# Prediction of zoonotic virus-host transmissibility using comparative airway organoids

**DOI:** 10.1101/2025.09.12.675569

**Authors:** Namwook Kim, Christine S. Kim, Junhee Park, Hyeon Jun Yoon, Ju Ang Kim, Yong Gun Kim, Joonhyung Kim, Injun Song, Donghyuck Ahn, Jihyeon Myeong, Byungmoo Oh, Jaeyoon You, Eunju Hong, Sukin Jeong, Kyungmoo Yea, Sung Won Kim, Ok Sarah Shin, Seung Joon Kim, Minho Lee, Myungin Baek, Youngtae Jeong

## Abstract

The emergence of new pandemic zoonotic diseases is accelerating. However, current approaches to control after outbreaks often allow the epidemic or pandemic spread of new zoonotic diseases, exemplified by the case of COVID-19. Prediction of zoonotic transmissibility will help minimize the outbreaks and socioeconomic damage. Here, we established highly efficient comparative airway organoids with minimal components from a panel of mammalian species for long-term culture and passaging and validated their utility to predict the transmissibility of zoonotic pathogens. Airway organoids from each species recapitulated the airway histology and contained the major cell types in the airway. They recapitulated the known susceptibility of their host species and tissues to diverse viruses. Transcriptomic analyses revealed that organoids from susceptible species induced genes of anti-viral immune responses only in common upon exposure to the same virus, although each species organoids displayed differential gene expression profiles. Also, organoids of the same species displayed differential transcriptomic responses against different viruses but induced anti-viral immune genes only in common. Our work provides a roadmap for comprehensive species- and tissue-level organoid panels for quick and reproducible uncharacterized pathogen surveillance to predict and prevent the next pandemic.

## 1 Introduction

Since the beginning of the 21^st^ century, consecutive outbreaks of pandemic or epidemic diseases, including COVID-19, SARS, swine flu (H1N1), Middle East Respiratory Syndrome, and Zika virus diseases, have posed severe threats to global human health. All of these global diseases originated from animal pathogens, and the emergence of new zoonotic diseases is accelerating. Current approaches for zoonotic disease control, including early isolation and the development of therapeutic drugs and vaccines, begin once a human outbreak is detected. During the lag time between the initial outbreak detection and its control, a new zoonosis could spread up to a pandemic level as exemplified by the case of COVID-19. Timely prediction of zoonotic transmissibility, identification of intermediate host animals, and early action for control could minimize new zoonotic outbreaks and spread.

Numerous efforts have been made to predict the emergence of new zoonosis, including metagenomic analyses, estimation of receptor specificity, infection of cell lines or tissues, and animal surveillance (1–3). However, these methods have limited accuracy in predicting real-world human or intermediate host infectivity. For example, although structural analysis predicted that SARS-CoV-2 binds to porcine ACE2, suggesting that the virus may infect pigs (4), direct inoculation of SARS-CoV-2 has failed to do so (5–7). Also, concerns have been raised that viral susceptibility testing in 2-dimensional (2D) cell culture does not faithfully recapitulate real *in vivo* infectivity, possibly due to the lack of organ microstructure and the loss of cell type diversity by dominant cell clones (1,8). Direct testing with tissue slices or in live organisms requires a large number of sometimes endangered wild animals, posing ethical and practical concerns. All these challenges emphasize the need for improved models to screen virus-host tropism.

Organoids are stem cell-derived 3D structures recapitulating the histology and gene expression pattern of originating tissues (9,10). Organoids can be passaged and have been successfully employed for modeling the infection of diverse pathogens (11–14). Indeed, previous studies reported the use of human airway organoids in testing and modeling the infection of diverse respiratory viruses, including influenza virus (12), respiratory syncytial virus (13), and SARS-CoV-2 (14–17). Furthermore, recent studies reported the establishment of bat organoids and their application of modeling SARS-CoV infection (18,19), suggesting the potential application of animal organoids in testing and modeling viral infection. Therefore, organoids from diverse animal species could overcome the limitations of current approaches and help predict zoonotic transmissibility.

Here, we report the establishment of airway organoids derived from five representative mammalian species to rapidly and accurately assess the tropism of zoonotic viruses. Infection of these organoids with respiratory viruses possessing differential host tropism provides proof-of-principle data that a panel of airway organoids from diverse animal species can provide a robust and accurate tool to predict cross-species transmissibility for predicting the next zoonotic pandemic.

## Methods and Materials

Detailed methods can be found in the supplementary material.

### Tissue Samples

**Mice:** All animal work complied with the Republic of Korea Animal Scientific Procedures Act and was approved by the Ethical Committee and the Institutional Animal Care and Use Committee of DGIST. C57BL/J specific-pathogen-free mice were obtained from Jackson laboratories, bred, and maintained in Daegu Gyeongbuk Institute for Science and Technology Animal Facility.

**Cow, pig, and dog:** Cow and pig tracheas were provided from eastern branch of Gyeongsangnam-do Veterinary Service (Yangsan, Korea). Dog tracheas were provided from Genia (Sungnam, Korea).

**Human tissues:** Human bronchi biopsy samples were obtained from patients who underwent surgery at Catholic University of Korea Seoul St. Mary’s Hospital, under the protocols approved by the institutional review board (Table S1). Human bronchial organoids were generated and used for this study under the protocols approved by the institutional review board at DGIST.

### Organoid generation

All tracheal or lung tissues were processed as previously described (20). Details are described in online methods.

### Virus infection

Organoids were transduced with viruses in serum-free medium for 2 h at MOI = 1. The source of viruses is listed in online methods.

## 2 Results and Discussion

### 2.1 Establishment of comparative airway organoids

The respiratory tract is one of the main routes for the infection and spread of contagious zoonotic pathogens. Therefore, we aimed to establish a panel of airway organoids from a set of mammalian species and validate its utility for predicting zoonotic virus-host tropism in the respiratory tract. To this end, we selected five animal species representing human, pest (mouse), pet (dog), and livestock (cow and pig) that are important sources or reservoirs of many zoonotic pathogens and also have close contact with humans (Figure 1A). We also selected three representative viruses well-known for their species tropism to examine as the proof-of-principle (Figure 1B).

**Figure 1.**
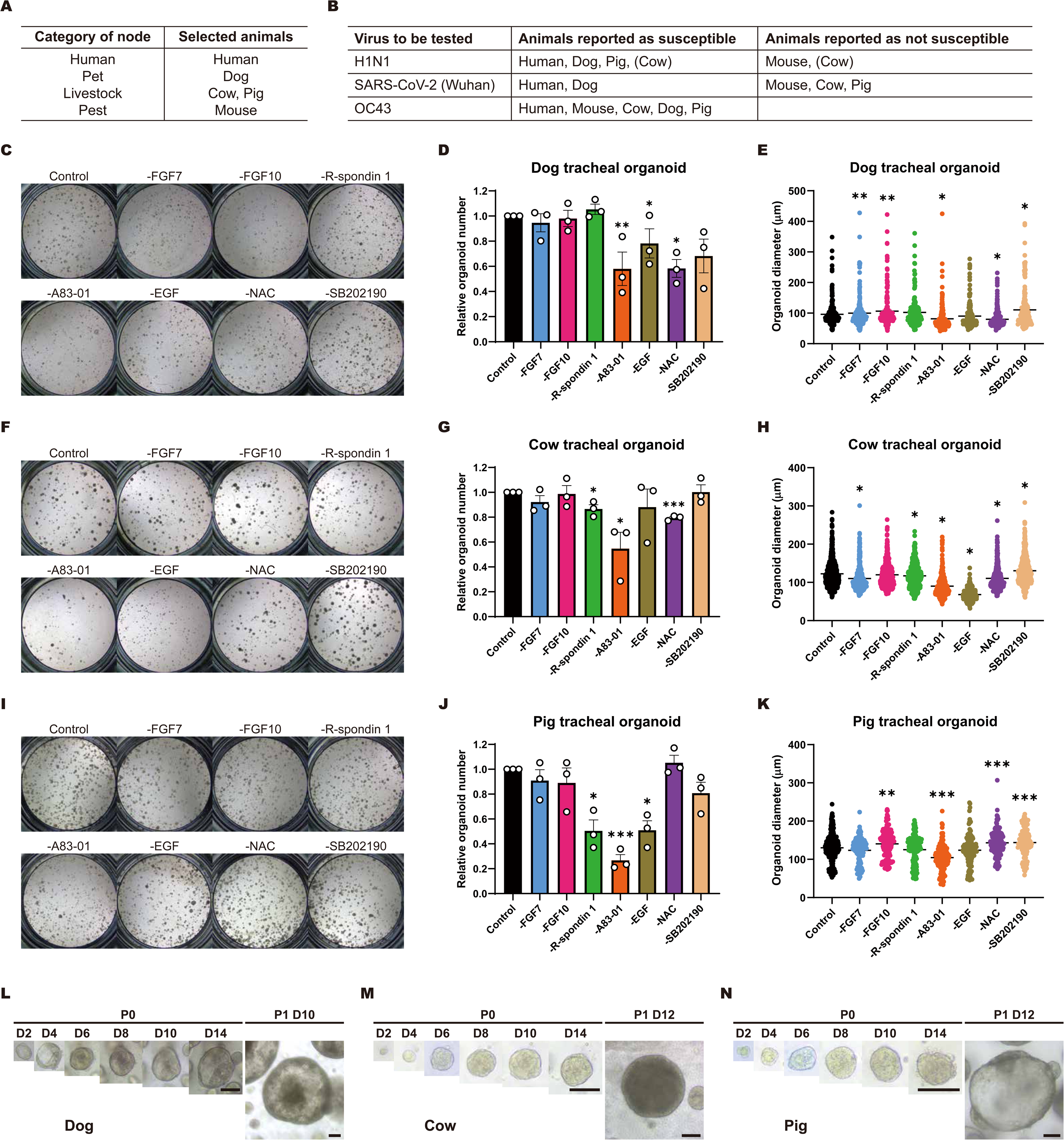
The establishment and optimization of long-term, three-dimensional cultures of airway organoids of dog, cow, and pig. A) Category of five representative animals selected in our platform in the zoonotic transmission B) Viruses tested in our platform as the proof-of-principle and their host susceptibility C-E) Optimization of dog airway organoid culture medium. Representative bright-field organoid images of each indicated culture condition (C). Relative numbers of organoids in each culture condition (D) and sizes of organoids in each culture condition (E). F-H) Optimization of cow airway organoid culture medium. Representative bright-field organoid images of each indicated culture condition (F). Relative numbers of organoids in each culture condition (G) and sizes of organoids in each culture condition (H). I-K) Optimization of pig airway organoid culture medium. Representative bright-field organoid images of each indicated culture condition (I). Relative numbers of organoids in each culture condition (J) and sizes of organoids in each culture condition (K). All data from c to k are collected from 3 biological replicates and presented as mean ± S.E.M (*, p < 0.05; **, p < 0.01; ***, p < 0.001). L) Time lapse images of dog airway organoids. M) Time lapse images of cow airway organoids. N) Time lapse images of pig airway organoids. Scale bars: 100 μm.

While previous studies reported the human and mouse airway organoid systems (20,21), the detailed culture conditions for airway organoids of dog and cow were not reported. Although a recent study reported the establishment of pig tracheal organoids, they used a commercial media for human airway organoids rather than optimizing a species-specific culture condition for pig tracheal organoids (22). Therefore, we first developed dog and cow airway organoids and optimized species-specific airway organoid culture conditions containing the minimal components necessary to efficiently generate dog, cow, and pig airway organoids displaying a pseudo-stratified epithelium-like histology, from primary single cells dissociated from dog, cow, and pig tracheas. To this end, we added or removed diverse factors starting from human airway organoid culture media (21) and customized the each species-specific airway organoid culture condition that allows the highest organoid-forming efficiency both in number and size. For the dog tracheal organoids, removal of A83-01, epidermal growth factor (EGF), or N-acetylcysteine (NAC) decreased either the number or size of the organoids, while removal of fibroblast growth factor 7 (FGF7), FGF10, and SB202190 rather increased the size of organoids. However, R-spondin-1 did not cause any significant change in the dog tracheal organoid formation (Figure 1C-E). For the cow tracheal organoids, removal of FGF7, R-spondin-1, A83-01, EGF, and NAC decreased either the number or size of the organoids, while removal of SB202190 increased the size of organoids (Figure 1F-H). On the other hand, FGF10 did not appear to have any significant effect in cow tracheal organoid formation. For pig tracheal organoids, removal of R-spondin-1, A83-01, and EGF decreased the number or size of the organoids, while removal of FGF10, NAC, and SB202190 increased the size of the organoids (Figure 1I-K). However, FGF7 influenced minimal effect in pig tracheal organoid formation. Additionally, we established human and mouse airway organoid cultures as previously described (21). When all airway organoids of each species were cultured by the optimized media, they reached the size of 100–500 μm in diameter within 10–14 days (Figure 1L-N and Figure S1).

We further examined whether these organoids displayed the expected histology of pseudo-stratified epithelium (Figure 2A-E and Figure S2). They contained basal cells (keratin 5 (*Krt5*)-positive), club cells (*Scgb1a1*-positive or polymeric immunoglobulin receptor (pIgR)-positive), and ciliated cells (*FoxJ1*-positive or acetylated tubulin (AcT)-positive), recapitulating the major cell types present in the tracheal and bronchial epithelium. While most primary airway organoids displayed cystic morphology, some organoids displayed solid morphology (Figure 2C).

**Figure 2.**
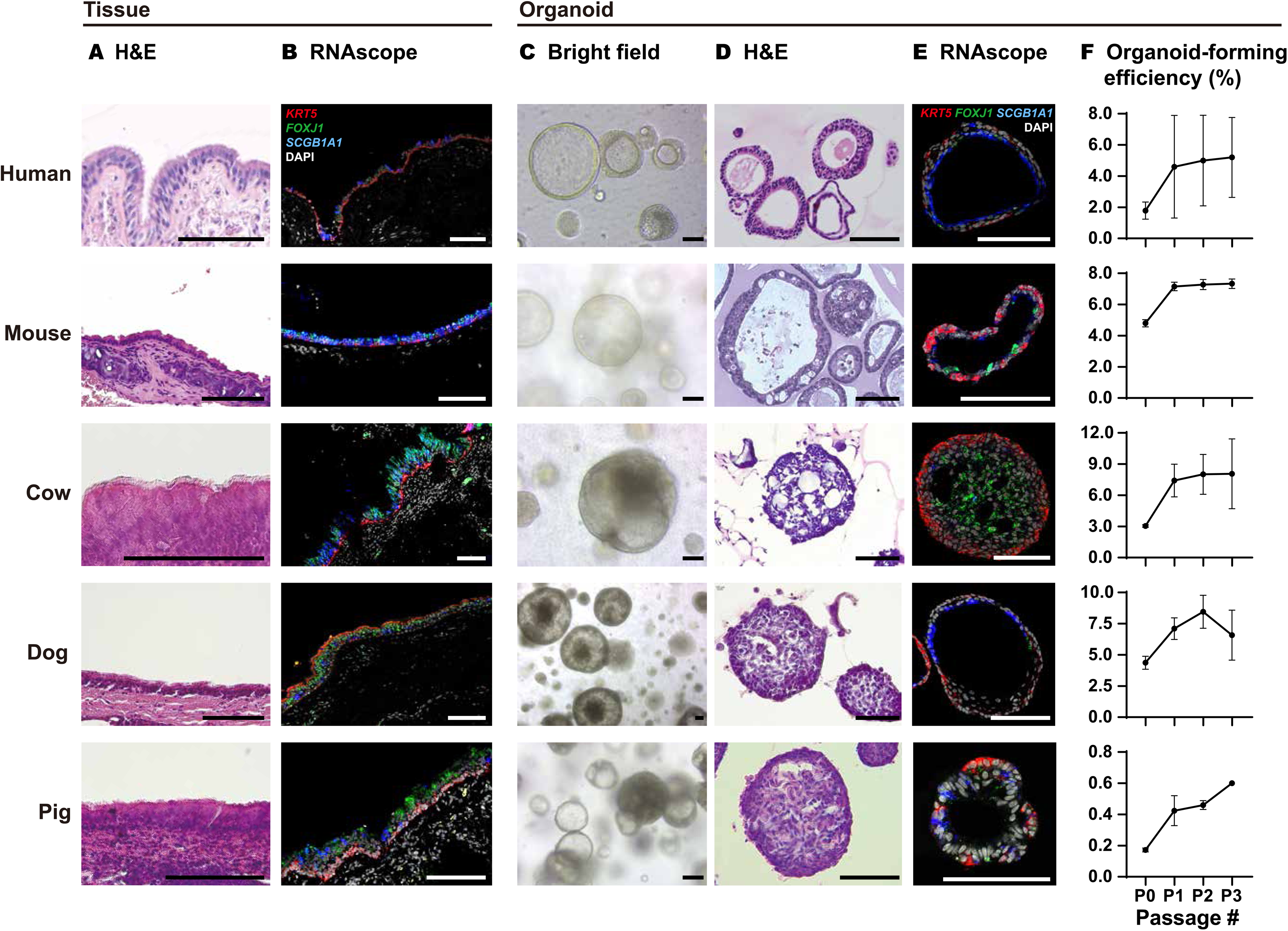
Organotypic representation of airway organoids of human, mouse, cow, dog, and pig. A) H&E staining of tracheal (mouse, cow, dog, and pig) or bronchial (human) epithelia. B) RNAscope images of tracheal or bronchial epithelia probed for *Krt5* (red), *Scgb1a1* (blue), and *Foxj1* (green). Images are single confocal planes in the middle of the organoids. C) Bright-field images of tracheal or bronchial organoids. D) H&E staining of tracheal or bronchial organoids. E) RNAscope images of tracheal or bronchial organoids probed for *Krt5* (red), *Scgb1a1* (blue), and *Foxj1* (green). Images are single confocal planes in the middle of the organoids. F) Organoid-forming efficiency of tracheal or bronchial organoids along passages (N=3). Scale bars: 100 μm.

All of airway organoids could be cultured and grown over 1–4 weeks at each passage and maintained for at least 6 passages over 6 months (Figure 1L-N and Figure S3). The organoid-forming efficiency of the primary airway cells reached to 2% in human, 5% in mouse, 4.8% in dog, 3% in cow, and 0.2% in pig (Figure 2F). While the organoid-forming efficiency increased over the first two passages, it reached to a plateau after the second or third passages. Notably, throughout the passaging process, the portion of solid organoids increased in all organoids. We therefore employed organoids only from primary tissue to passage number 3 for testing viral infectivity.

In sum, we optimized airway organoid culture conditions for diverse animal species, in addition to previously reported human and mouse airway organoid cultures, thereby establishing a panel of comparative airway organoids that recapitulates the original tissue histology and can be maintained *in vitro* over long periods of time.

### 2.2 Differential infection of airway organoids by human influenza A virus (H1N1)

We next examined whether these airway organoids mirror their source organisms with respect to virus-host organismal tropism. To this end, we infected these organoids with multiple respiratory viruses that are known to have different species tropism (Figure 1B). We first tested H1N1 (A/human/Korea/KUMC-33/2006/H1N1), a strain of human influenza A virus (IAV) that is relatively well characterized for its species tropism and its transmissibility from humans to animals and *vice versa*. Interspecies transmission of IAV between humans and pigs (23–25) or opportunistically to dogs (26,27) has been reported. However, human influenza strains do not attach to murine airway or alveoli cells (28,29) since mice express α2,3-linked sialic acid (SA) instead of α2,6-linked SA, the receptor for human IAV. In the case of cows, serum and nasal mucus analyses have indicated the presence of antibodies against human H1N1 (30,31). Also, a recent study reported that α2,6-linked SA is minimally expressed in bovine trachea but significantly expressed in bovine lung (32), suggesting that although cows could be susceptible to human H1N1, it would infect and replicate less efficiently in cow tracheal organoids.

We therefore hypothesized that the H1N1 virus would infect and replicate in the airway organoids of human, dog, and pig, but not those of mouse and cow. To test our hypothesis, we first examined whether organoids recapitulated the pattern of tissue expression of α2,6-linked SA. As reported in the airway tissues, human, dog, and cow airway organoids expressed significantly high levels of α2,6-linked SA (Figure 3A-B). However, mouse and cow airway organoids minimally expressed α2,6-linked SA.

**Figure 3.**
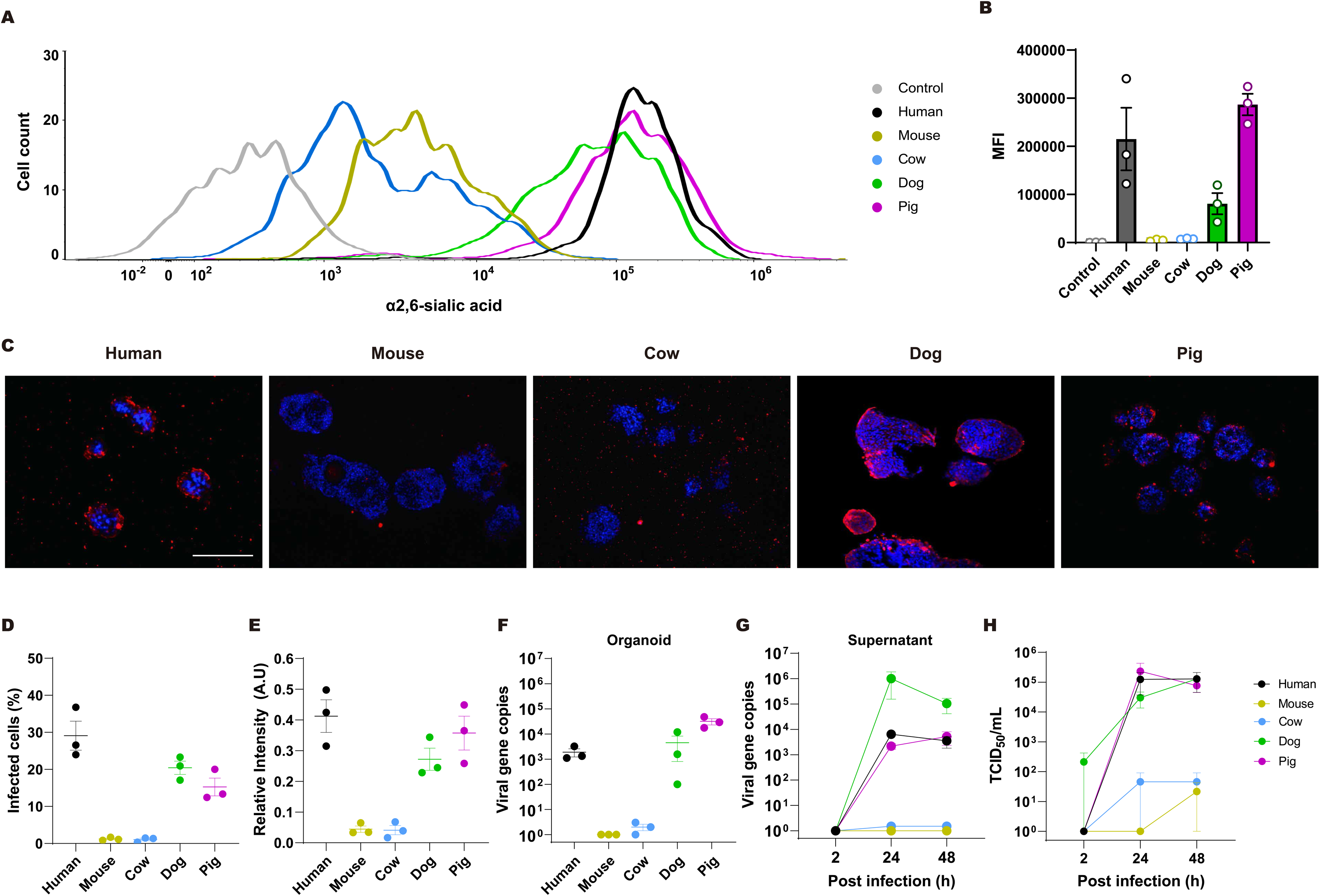
Investigation of H1N1 viral tropism in airway organoids of five mammalian species. A-B) Expression of of α2,6-linked SA in animal airway organoids in our panel. Representative graph analyzed by flow cytometry (A) and mean fluorescence intensity (MFI) (B) are shown (N=3). C) Immunofluorescence staining of H1N1 nucleoprotein in airway organoids at 48 hpi of H1N1 at MOI = 1. Representative images are shown from N=3. Scale bars: 100 μm. D) The percentage of viral nucleoprotein-expressing cells over total cells in airway organoids. E) Relative fluorescence signal intensity of H1N1 nucleoprotein over H1N1-infected airway organoids. A.U.: arbitrary unit. F) Quantification of viral gene copy numbers per μg of total RNA in airway organoids infected with H1N1 virus. Total viral gene copy numbers were measured by qPCR. G) Quantification of viral gene copy numbers per mL of culture supernatants of airway organoids. Cell-free media were harvested at 2, 24, and 48 hpi, and viral gene copy numbers were quantified by qPCR. H) Viral titers in the organoid culture supernatants were determined by 50% tissue culture infectious dose (TCID_50_) assay at the indicated time points. All experiments were done with three biological replicates.

Next, we inoculated H1N1 into the organoid cultures at a multiplicity of infection (MOI) of 1. Then, we measured the percentage of infected cells by immunostaining of H1N1 nucleoprotein (NP) in organoids and quantified the viral gene copy number by qPCR in both organoids and their culture supernatants. About 10-30% of cells in human, dog, and pig airway organoids expressed NP, while the percentage of NP-positive cells in cow and mouse organoids was negligible (Figure 3C-E). Consistently, between 24 and 48 hrs post-infection (hpi), the viral gene copy number was more than 1,000-fold increased both in the organoids and in the culture media of human, dog, and pig organoids, while negligible in the organoids and in the culture media of mouse and cow tracheal organoids (Figure 3F-G). Furthermore, human, dog, and pig organoids produced significantly increased amount of replicable H1N1 viruses when they were infected, while mouse and cow organoids released minimal amount (Figure 3H).

These data indicate that comparative airway organoids recapitulate the known species tropism of H1N1 virus.

### 2.3 Differential infection of cow airway and bronchioalveolar organoids by H1N1 representing tissue tropism

Recently, an outbreak of highly pathogenic H5N1 avian influenza in dairy cows was reported (33). Subsequent microscopic analysis revealed that H5N1 can infect the alveolar cells of cows (34,35). Since lung cells of cows express α2,6-linked SA, in contrast to tracheal cells (32), we further examined whether H1N1 could infect cow lung organoids, whereas it failed to infect cow tracheal organoids, as shown in Figure 3.

To this end, we sought to establish cow lung organoids. Organoids from cow lung tissue grew up to the size of 300 μm in diameter within 10–14 days when cultured with human airway organoid culture media supplemented with Insulin-Transferrin-Selenium (ITS), which allowed long-term culture and passaging for at least 6 passages over 6 months (Figure 4A and Figure S4A-C). The organoid-forming efficiency of primary cow lung cells reached 2.5% and increased over the first two passages (Figure S4D). They exhibited cystic or alveoli-like morphology and contained SPC+ alveolar type II cells and AQP5+ alveolar type I cells and expressed significantly higher levels of lung marker genes (*Nkx2-1*, *Sftpc*, and *Ager*) than airway organoids (Figure 4B-G). However, they also expressed *Krt5*, *Scgb1a1*, and *Foxj1* at modest levels, suggesting that these organoids are bronchioalveolar organoids rather than alveolar organoids (Figure 4G). Nevertheless, we could still employ the cow bronchioalveolar organoids to test whether H1N1 can infect cow lungs, as these organoids contain both alveolar type I and type II cells, which are found in alveolar epithelia. Indeed, in contrast to the cow tracheal organoids, cow bronchioalveolar organoids expressed α2,6-linked SA at a significantly higher level (Figure 4H-I) and also exhibited positive signals against H1N1 NP when inoculated with human H1N1 (Figure 4J-L). Furthermore, the viral gene copy number increased more than 10,000-fold in the organoids and in the organoid culture supernatants (Figure 4M-N). In addition, cow lung organoids could release infectious virus to the supernatant (Figure 4O). Considering that H1N1 infects cow bronchioalveolar organoids but not cow tracheal organoids, these data indicate that comparative airway organoids recapitulate the virus-tissue tropism as well as the virus-host tropism.

**Figure 4.**
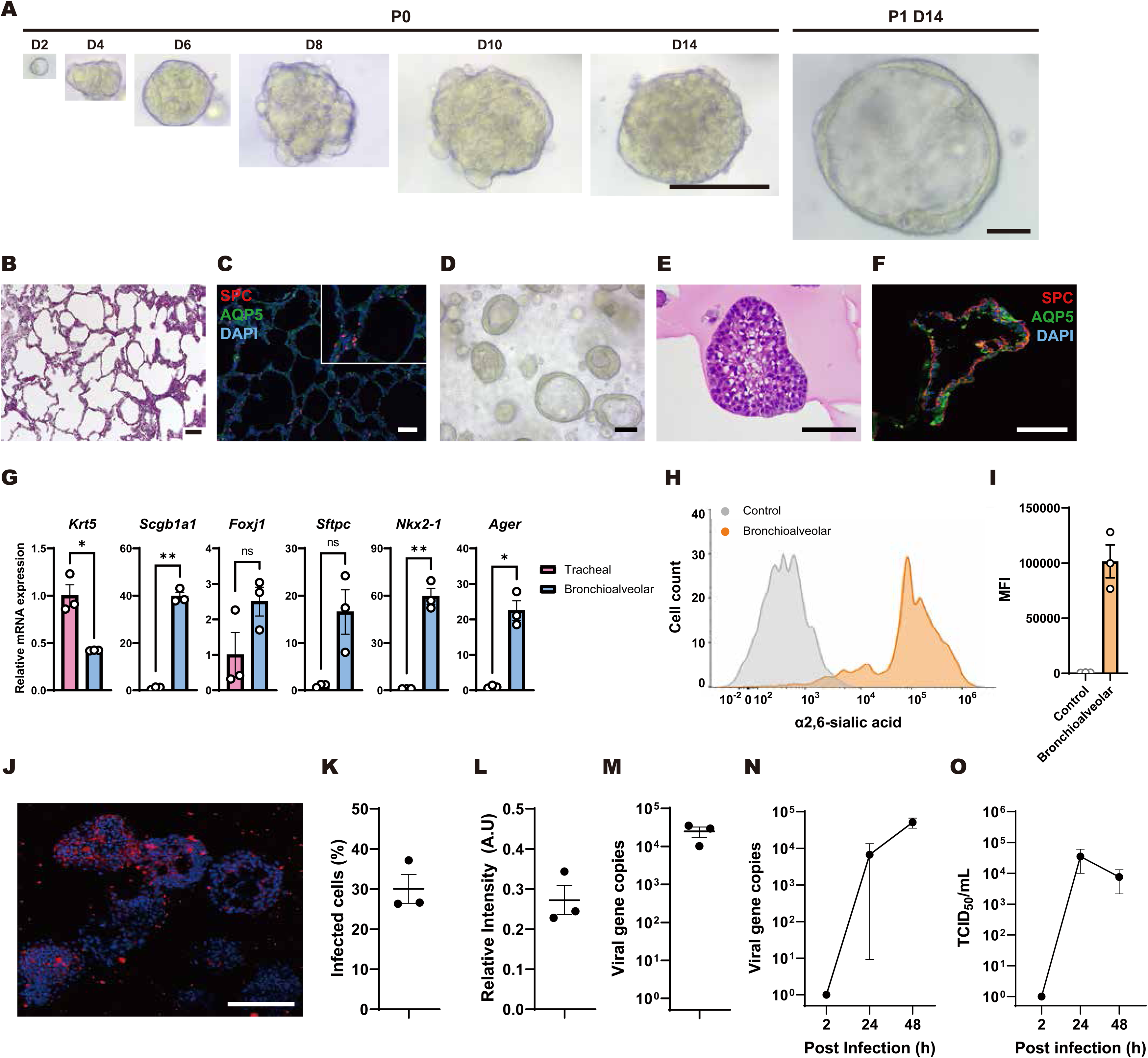
Establishment of cow bronchioalveolar organoids and investigation of H1N1 viral infectivity. A) Time lapse images of cow bronchioalveolar organoids. Scale bars: 100 μm. B) H&E staining of cow bronchioalveolar epithelia. C) Immunofluorescence staining of cow bronchioalveolar epithelium. AQP5 and SPC were used as the markers for alveolar type I and II cells, respectively. D) Bright-field images of cow bronchioalveolar organoids. E) H&E staining of cow bronchioalveolar organoids. F) Immunofluorescence staining of cow bronchioalveolar organoids. G) Relative gene expression of representative lung and tracheal markers in organoids generated from cow lung and tracheal tissues assessed by qPCR (N=3). All data are presented as mean ± S.E.M (*, p < 0.05; **, p < 0.01; ns, not significant). H-I) Expression of of α2,6-linked SA in cow lung organoids. Representative graph analyzed by flow cytometry (H) and mean fluorescence intensity (MFI) (I) are shown (N=3). J) Immunofluorescence staining of H1N1 nucleoprotein in cow bronchioalveolar organoids at 48 hpi of H1N1 at MOI = 1. Representative images are shown. K) The percentage of viral nucleoprotein-expressing cells over total cells in cow bronchioalveolar organoids. L) Relative fluorescence signal intensity of H1N1 nucleoprotein over H1N1-infected cow bronchioalveolar organoids. A.U.: arbitrary unit. M) Quantification of viral gene copy numbers per μg of total RNA in cow bronchioalveolar organoids infected with H1N1 virus. Total viral gene copy numbers were measured by qPCR. N) Quantification of viral gene copy numbers per mL of culture supernatants of cow bronchioalveolar organoids. Cell-free media were harvested at 2, 24, and 48 hpi, and viral gene copy numbers were quantified by qPCR. O) Viral titers in the organoid culture supernatants were determined by 50% tissue culture infectious dose (TCID50) assay at the indicated time points. All experiments were done with three biological replicates.

### 2.4 Differential infection of airway organoids by SARS-CoV-2

To further test whether organoids reflect the species tropism of viral infection, we next explored the infectivity of SARS-CoV-2 that evoked pandemic COVID-19. Although the pseudo-entry SARS-CoV-2 is a pseudotyped lentivirus and not a real SARS-CoV-2, it expresses the spike protein of SARS-CoV-2 and, therefore, provides a proxy for the infectivity of SARS-CoV-2 (36). It is documented that the spike protein of SARS-CoV-2 binds to human ACE2 but does not bind to murine ACE2 (4), having standard experimental mice uninfected by SARS-CoV-2. Direct intranasal or intratracheal inoculation of SARS-CoV-2 did not induce clinical signs or viral replication in pigs (5–7). In contrast, studies reported the susceptibility of dogs to SARS-CoV-2, raising a concern that pets could become intermediate reservoirs for SARS-CoV-2 (37,38). Although Wuhan-like SARS-CoV-2 isolates induced seroreactivity from one-third of inoculated cattle, it did not induce productive viral replication or infection (39–41). Therefore, we predicted that pseudo-entry SARS-CoV-2 would infect human and dog organoids but not mouse, cow, and pig airway organoids. As predicted, 8-12% of cells in human and dog airway organoids displayed positive signals upon immunostaining against spike protein, while airway organoids of mouse, cow, and pig were refractory to infection (Figure 5A-C). Since pseudo-entry virus can infect but not replicate in cells, we did not measure the viral gene copy number in organoids or culture supernatants. Nonetheless, these data support that comparative airway organoids recapitulate the known species tropism of SARS-CoV-2.

**Figure 5.**
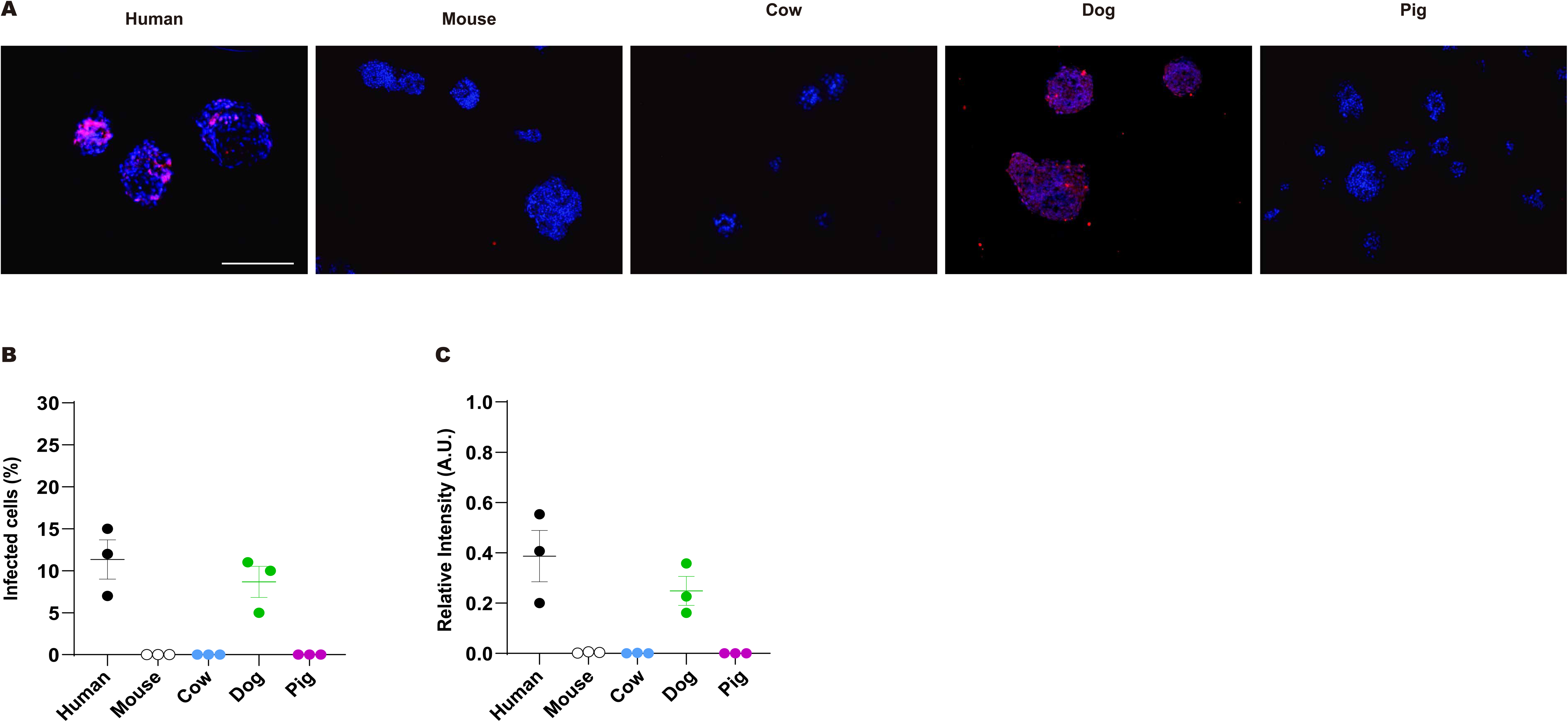
Differential ability of pseudo-entry SARS-CoV-2 to infect the multi-species airway organoid panel. A) Immunofluorescence staining of spike protein in airway organoids at 48 hpi of pseudo-entry SARS-CoV-2 at MOI = 1. Representative images are shown from N=3. Scale bars: 100 μm. B) The percentage of SARS-CoV-2 spike protein-expressing cells over total cells in airway organoids. C) Relative fluorescence signal intensity of spike protein over pseudo-entry SARS-CoV-2-infected airway organoids. A.U.: arbitrary unit. All experiments were done with three biological replicates.

### 2.5 Differential infection of airway organoids by seasonal coronavirus

OC43 is another member of the betacoronavirus family and causes seasonal common cold in humans. OC43 can replicate in diverse murine organs including lung upon intranasal inoculation (42). Viral sequence and serologic analyses suggest that OC43 may infect dogs and pigs (43,44). Cows are accepted as an intermediate host based on high antigenic and genetic sequence similarities (45–47). Thus, in contrast to H1N1 and pseudo-entry SARS-CoV-2, which infected a subset of our animal organoids, we predicted that OC43 would infect all organoids in our panel. Indeed, OC43 not only expressed viral antigen in all organoids (Figure 6A-C) but also replicated actively in all organoids and supernatants in our panel (Figure 6D-E). These data further corroborate our conclusions that organoids mimic organism-level viral tropism across diverse respiratory pathogens.

**Figure 6.**
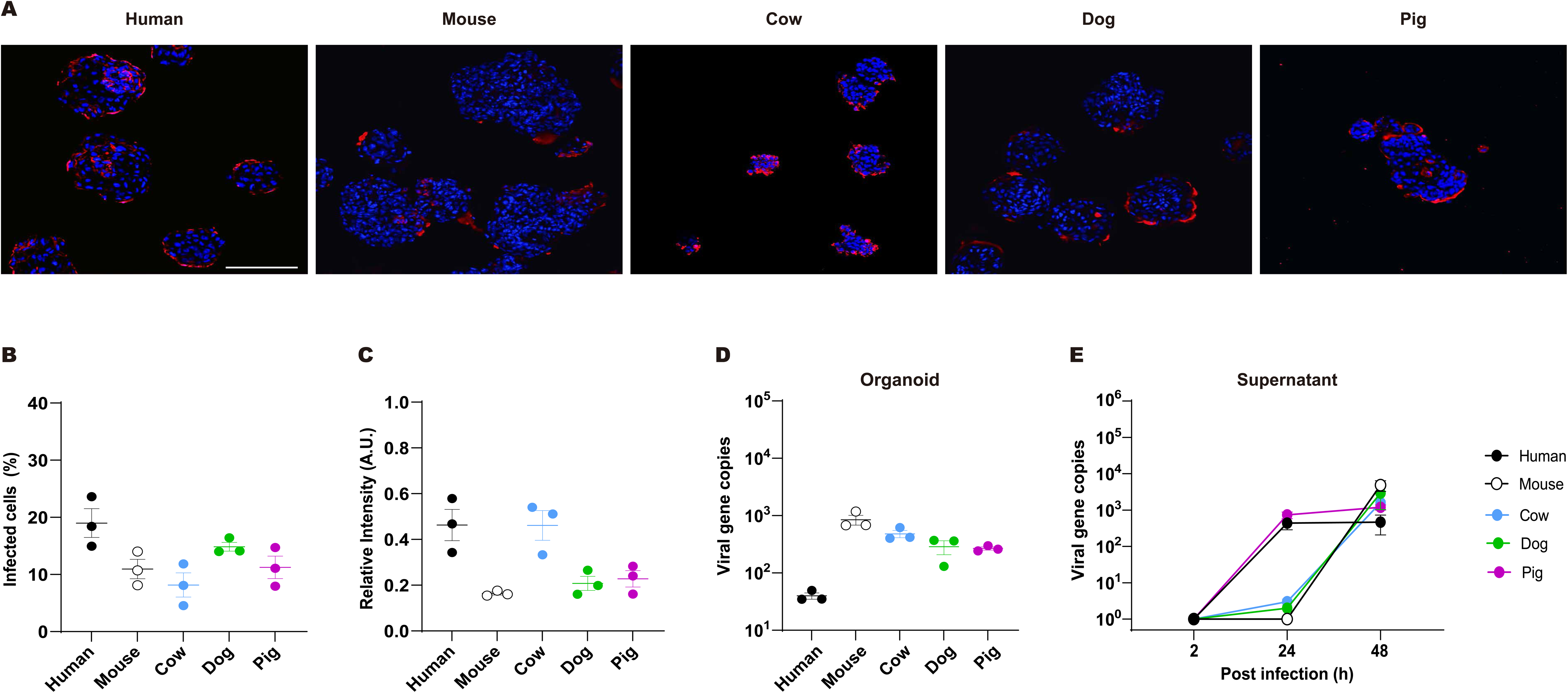
Profiling the infectivity of the OC43 betacoronavirus across five mammalian species using airway organoids. A) Immunofluorescence staining of nucleocapsid protein in airway organoids at 48 hpi of OC43 at MOI = 1. Representative images are shown from N=3. Scale bars: 100 μm. B) The percentage of nucleocapsid protein-expressing cells over total cells in airway organoids. C) Relative fluorescence signal intensity of nucleoprotein over OC43-infected airway organoids. A.U.: arbitrary unit. D) Quantification of viral gene copy numbers per μg of total RNA in airway organoids infected with OC43 virus. Total viral gene copy numbers were measured by qPCR. E) Quantification of viral gene copy numbers per mL of culture supernatants of airway organoids. Cell-free media were harvested at 2, 24, and 48 hpi, and viral gene copy numbers were quantified by qPCR. All experiments were done with three biological replicates.

### 2.6 Transcriptome analysis of tropic and non-tropic viral responses

Previous studies showed that organoids display similar transcriptomic responses to the corresponding patient organs after viral infection (15,48). Therefore, we next aimed to identify any distinctive virus-induced genetic responses shared across species only by the virus-tropic species organoid pairs, which can serve as a potential biomarker of zoonotic viral transmissibility. Viral infection elicits antiviral innate immune responses, including the production of type I interferons and inflammatory cytokines (49). We therefore evaluated the changes in the expression of 20 representative genes related to the antiviral and innate immune pathways in response to H1N1 infection in airway organoids from tropic (human and dog) and non-tropic (mouse and cow) species. Intriguingly, H1N1 infection significantly induced the expression of antiviral and innate immune pathway-related genes in human and dog airway organoids, but not in mouse and cow airway organoids at 2 days after viral inoculation (Figure 7A).

**Figure 7.**
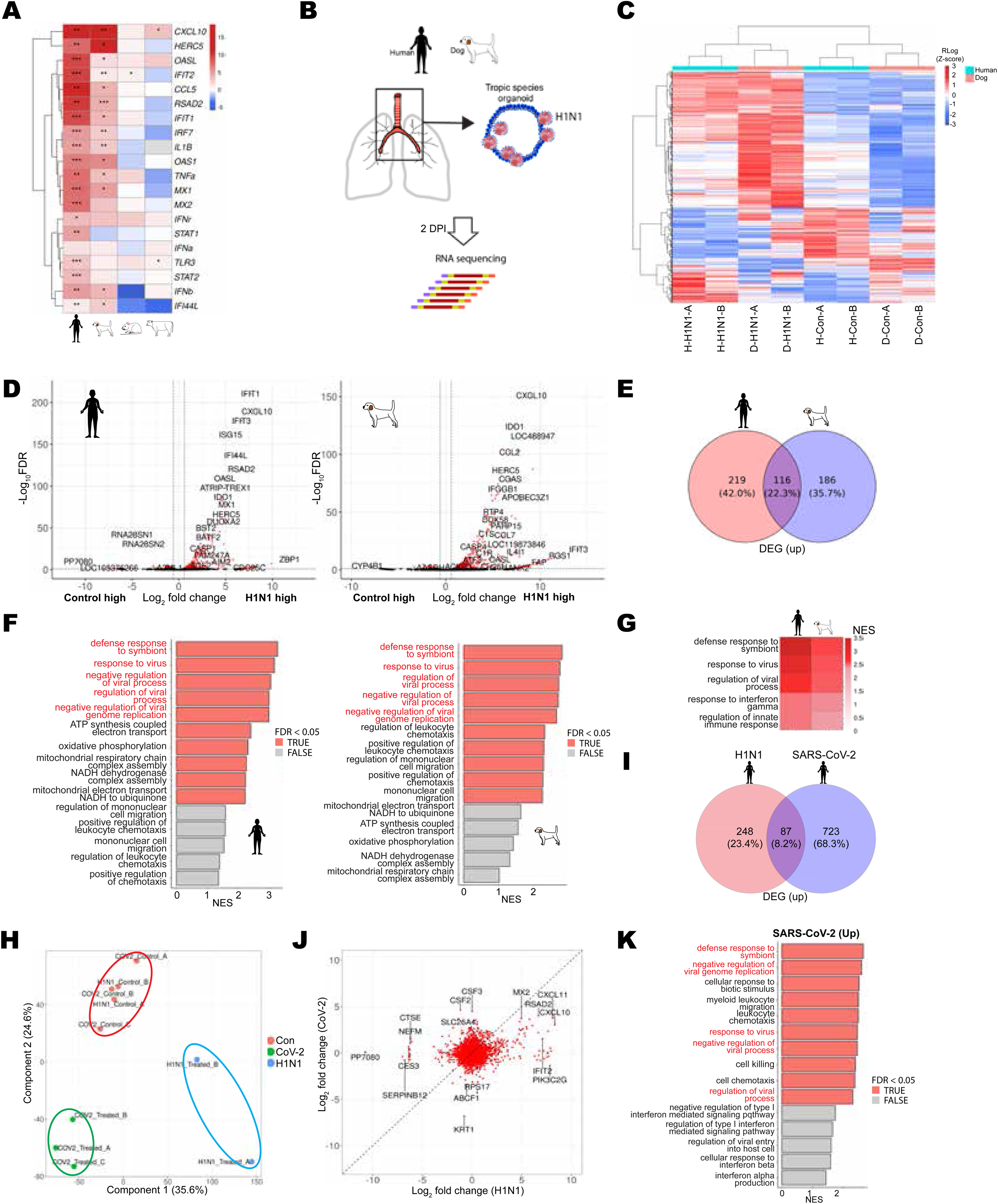
Shared transcriptomic responses of viral infection across host species and across viral pathogens. B) Heat map showing the gene expression changes of the representative 20 genes in the interferon or innate immune pathways compared between H1N1-infected and mock-infected organoids in each species. Gene expression was measured by qPCR (*, p < 0.05; **, p < 0.01; ***, p < 0.001). C) Schematic representation of experimental design for RNA sequencing. D) Unsupervised hierarchically clustered heatmap of 614 DEGs from each species’ organoids with or without H1N1 viral infection. H: human, D: dog. E) Volcano plot of DEGs in H1N1-infected human and dog airway organoids. DEGs with FDR < 0.05 and log2FC (Fold Change) > log_2_(1.5) are indicated. Each dot represents individual genes and denoted by gene name. F) Venn diagram of DEGs in H1N1-infected human and dog airway organoids. G) Bar graph representing GSEA for GO:BP upregulated in H1N1-infected human and dog airway organoids compared to mock-infected organoids. H) Heat map representing the GSEA of the top 5 upregulated gene sets shared by human and dog airway organoids from H1N1-infected organoids compared to mock-infected organoids. I) Principal component analysis of differential gene expression from H1N1-or SARS-CoV-2-infected human airway organoids compared to mock-infected organoids. J) Venn diagram of DEGs in H1N1-or SARS-CoV-2-infected human airway organoids. K) Scatter plot of whole transcripts in H1N1-or SARS-CoV-2-infected human airway organoids. Key DEGs are denoted by gene name. L) Bar graph representing GSEA upregulated in SARS-CoV-2-infected human airway organoids compared to mock-infected organoids.

To analyze gene expression changes in an unbiased manner, we performed bulk RNA sequencing on human and dog airway organoids at 2 days after H1N1 inoculation. This allowed us to identify differentially expressed genes (DEGs) shared by organoids from species susceptible to H1N1 (Figure 7B and Table S2). Although H1N1 treatment induced distinct changes in gene expression between human and dog organoids (Figure 7C-D), more than 20% of DEGs were shared in both human and dog airway organoids (Figure 7E and Table S3). To identify the genetic signatures of H1N1 infection, we first performed gene set enrichment analysis (GSEA) on transcriptome profiles induced by H1N1 administration in airway organoids from each species separately. Both species-specific and shared gene sets were identified. Among the species-specific gene sets, those related to oxidative phosphorylation and mitochondrial respiration were enriched in human bronchial organoids, while gene sets related to leukocyte chemotaxis and migration were enriched in dog tracheal organoids (Figure 7F and Figure S5A). Of note, gene sets such as response to virus, response to interferon gamma, and regulation of innate immune response, all of which are related to antiviral innate immune responses, were induced by H1N1 both in human and dog organoids (Figure 7F-G).

Similar results were derived when we identified shared genes and species-specific DEGs first, and then performed GSEA of shared genes and species-specific DEGs. While gene sets related to pyroptosis and NF-kappaB signal transduction were specifically enriched in human airway organoids, gene sets related to the phosphorylation of STAT proteins and immune cell proliferation were specifically enriched in dog airway organoids (Figure S5A). However, GSEA of 116 shared genes induced by H1N1 both in human and dog organoids revealed the enrichment of gene sets related to response to virus and viral replication. These findings underscore that antiviral innate responses are common cellular responses induced by H1N1 across tropic host species.

Following our finding that H1N1 viral infection induces genes related to antiviral innate immune responses in airway organoids across tropic species, we next explored how organoids of the same species respond to different viruses. To this end, we compared transcriptome profiles of our H1N1 dataset with a publicly available transcriptome dataset (GSE150819) from SARS-CoV-2-treated human bronchial organoids. Principal component analysis and unsupervised hierarchical clustering clearly separated H1N1-infected human bronchial organoids from SARS-CoV-2-infected human bronchial organoids (Figure 7H and Figure S5B). Only 90 (4.69%) genes out of 1,916 DEGs were commonly up-or down-regulated by H1N1 and SARS-CoV-2 infection (Figure 7I-J, Figure S5C, D, and F, and Table S4), emphasizing the distinct cellular responses to these viruses. Of note, gene sets related to ’defense response to virus’ and ’defense response to symbiont’ were the only commonly enriched gene sets (Figure 7K and S5E-F).

Taken together, these data show that antiviral innate immune responses are the only shared responses elicited in airway organoids from tropic animals, and these responses are not specific to particular viral species. These data further indicate that transcriptomic changes of antiviral innate immune responses, together with organoids, can serve as a biomarker for viral infection or tropism, further underscoring the utility of the comparative organoid platform as an improved proxy predicting zoonotic transmissibility.

## Discussion

In this study, we report that comparative airway organoids can be used as an improved platform to predict the cross-species transmissibility of emerging zoonotic viruses (Figure S6). Establishing comprehensive respiratory organoids from diverse categories of animals and applying a set of respiratory viruses with different species tropism, we demonstrate that the airway organoids faithfully recapitulate known organismal and tissue infection patterns, providing a significantly improved platform to screen the transmissibility of emerging zoonotic viruses. Furthermore, we show that transcriptomic changes of antiviral and innate immune genes are induced by viral infection, irrespective of the viral or host species when host species is susceptible to the viral infection. This proposes their potential utility, in combination with organoids, as a biomarker in predicting zoonotic transmissibility.

To establish comparative organoids that can be used to predict cross-species pathogen transmissibility, we first optimized airway organoid culture conditions individually customized for each species of animals reported as important nodes (dog, cow, and pig) for zoonosis. Still, only a few organoid systems for animals other than human and mouse have been reported despite their important roles as the intermediate hosts and reservoirs in zoonotic diseases. Species-specific differences partly hampered the establishment of animal organoid systems since different species have different requirement of culture components. To address these limitations in the field, we established detailed highly efficient species-specific airway organoid culture conditions for dog, cow, and pig. Despite a recent report about an establishment of pig airway organoids, they utilized commercial human airway epithelial cell culture medium rather than optimizing culture condition for the pig airway organoids (22). In contrast, we optimized highly efficient culture conditions with minimal components specifically customized for each species, which allow long-term culture and passaging for all of our animal organoids at least more than 6 passages over 6 months.

Our comparative organoid platform provides a basis for the fundamentally different control system for emerging zoonotic diseases from ‘control after outbreak’ to ‘prediction and prevention before outbreak’. Current control system begins once a human outbreak is detected. During the time gap between initial outbreak and their control, new zoonoses could spread even to a pandemic level as exemplified by COVID-19 case. In contrast, comparative organoids enable to predict the cross-species transmissibility of diverse emerging pathogens and their variants and set up the development of vaccines and therapeutic drugs in a proactive way prior to a real-world human outbreak. Although diverse methods have been tested to predict zoonotic transmissibility, they had not been widely used due to their limited accuracy. For example, SARS-CoV-2 fails to infect pigs despite the multiple trials through intravenous, intratracheal, and intramuscular injections (5–7), although it is predicted to bind to porcine ACE2 in the structural analysis (4). In contrast, 3D organoids accurately recapitulated the virus-host species tropism as previously reported (Figure 3-6), indicating their utility in predicting the real viral infectivity. Although 2D-cultured human cells are sometimes used for testing the viral susceptibility, organoids have advantages in that organoids maintain the tissue microstructures and include all of the major cell types in the tissue, while a few cell clones become dominant in 2D-cultured cells (9,10).

In addition to identifying human viral infectibility, identifying and distancing from the intermediate hosts and reservoir animals is indispensable to prevent or effectively control zoonotic diseases. Indeed, recent studies reported that pets such as dogs and cats (37,38), livestock such as cows (39,41), and wild animals such as tigers, pangolins, and deer (50–53) can be infected by SARS-CoV-2 or its variants, raising the issue of intermediate hosts and reservoirs in the control of COVID-19. However, it is practically and ethically challenging to directly test many of these species, particularly endangered wild species. Utilization of the comparative organoids could diminish the need for full animal testing as a first line of inquiry and significantly increases the speed and throughput.

Our study further shows that multi-organ organoids can provide information on tissue tropism. The route of infection is a key piece of information to understand viral pathogenesis and establish preventive or therapeutic strategies. In our study, H1N1 infected cow lung organoids but did not infect cow tracheal organoids (Figure 3 and 4). In line with our data, a previous study reported the potential utility of organoids for screening SARS-CoV-2 infectivity in human tissues (54), although liver organoids were the only organoids included in that study.

Remarkably, our transcriptome analyses suggest that innate immune and inflammatory responses (55) are commonly activated in tropic host species or tissues. Furthermore, these gene sets related to ’antiviral innate immune responses’ are the only shared gene sets induced across host species and also across viral pathogens (Figure 7). Previous studies corroborated that viral infection typically induced genes in these biological pathways (14,15,54–57). By comparing the virus-induced transcriptomic responses across species, we further show that different host species induce these genes related to ’antiviral immune responses’ in common although each species has its own species-specific transcriptomic response to the same virus infection. Also, we show that these transcriptomic changes occur only to tropic species after viral infection, which indicated that gene expression changes related to ’antiviral innate immune responses’ are not evoked by simple viral encounter. Therefore, our study suggests transcriptomic changes in genes related to ’antiviral immune responses’ as a biomarker differentiating viral tropism across species.

The major limitation of our study is that although we demonstrated the utility of comparative organoids in predicting virus-host tropism, we employed only respiratory organoids. However, the route of infection and virus-tissue tropism are also key pieces of information in determining the virus-host tropism and to understand viral pathogenesis and establish preventive or therapeutic strategies. Therefore, we speculate that, expanding our study, establishment comparative, multi-organ organoid platform will provide more complete information on viral tropism to screen inter-species transmissibility of emerging zoonotic pathogens. In addition, we employed pseudo-entry SARS-CoV-2 due to the limited access to the biosafety level 3 facility. However, pseudo-typed viruses have been used as surrogates of the native viruses including SARS-CoV-2 and HKU5-CoV-2 for testing their infectivity (54,58–61). Additionally, although organoids composed of airway epithelial cells are sufficient in testing virus-host tropism, complex organoids including other tissue cells such as fibroblasts and immune cells could provide more comprehensive understanding of disease pathogenesis. Recently, organoids with immune cells or lymphoid tissues have been established and shown that they can mimic the humoral immune responses to influenza and rabies viruses (62,63). Although we did not include immune cells in our organoid system, airway organoids with lymphoid systems could be used for and facilitate the development and testing of vaccines for airway viruses since they can simulate human immune responses more physiologically accurately than animal models.

In conclusion, we establish comparative airway organoids and provide proof-of-principle data demonstrating that comparative airway organoids are an improved platform to test virus-host species tropism and tissue tropism. The extension of our organoid platform to a more comprehensive list of species and organ types will provide a critical resource for the prediction of zoonotic transmissibility and identification of tissue specificity of emerging viruses. Furthermore, we believe that our platform will ultimately revolutionize our zoonotic disease control system from ‘control after outbreak’ to ‘prediction and prevention before outbreak’ and help minimize the occurrence of new pandemic zoonotic diseases.

## Acknowledgement

We thank Ms. Jayoung Kim, a manager at Eastern Branch of Gyeongsangnam-do Veterinary Service and Jungi Lee, the deputy director at Jungbu Branch Office of Gyeongsangnam-do Regional Headquarter of Livestock Health Control Association for providing cow and pig trachea specimens. Also, we thank Korea Bank for Pathogenic Viruses for providing H1N1.

## Author Contributions

**Conception and design:** Y. Jeong, M. Baek

**Performing experiments**: N. Kim, C. Kim, J. Park, Y. G. Kim, I. Song, D. Ahn, B. Oh, J. Myeong, S. W. Kim, O. Shin, M. Baek, Y. Jeong

**Analysis and interpretation of data**: N. Kim, C. Kim, H. Yoon, J. Kim, J. Yoon, E. Hong, S. Jeong, K. Yea, S. J. Kim, M. Lee, M. Baek, Y. Jeong

**Writing, review, and/or revision of the manuscript**: M. Lee, M. Baek, Y. Jeong

