## Supplementary file for "Prediction of zoonotic virus-host transmissibility using comparative airway organoids"

### METHOD DETAILS

#### Experimental model and subject details

#### Mice

All experimental work complied with the Republic of Korea Animal Scientific Procedures Act. The animal study was approved by the Ethical Committee and the Institutional Animal Care and Use Committee of DGIST. C57BL/6J specific-pathogen-free (SPF) mice were obtained from Jackson laboratories, bred, and maintained in Daegu Gyeongbuk Institute of Science and Technology Animal Facility. 6-24 weeks old mice were used in this study.

#### Cows and pigs

Approximately 1x1x1 cm sized pieces of about 3 years old cow and about 6 months old pig tracheal or lung samples were obtained from the eastern branch of Gyeongsangnam-do Veterinary Service (Yangsan, Korea).

#### Dogs

Tracheal samples from about 30 months old beagle dogs were obtained from Genia (Sungnam, Korea).

#### Human tissues

Deidentified human bronchial biopsy samples with no background pathologies were obtained from lung cancer patients who underwent surgical treatment at the Seoul St. Mary's Hospital of the Catholic University of Korea, with written informed consent from donors under the protocols approved by the institutional review board. Overall, bronchial tissues from a total 10 donors were used in this study to set up human airway organoid culture, and randomly picked three organoid lines were used to test viral infectivity (supplementary table 1).

**Supplementary table 1. Clinical information of tissue donors.**

| Patient ID | Hospital | Age | Sex | Smoking status | Tissue provided |
|---|---|---|---|---|---|
| 1 | Seoul St. Mary's Hospital | 65 | Female | Current | Bronchus |
| 2 | Seoul St. Mary's Hospital | 68 | Female | Never | Bronchus |
| 3 | Seoul St. Mary's Hospital | 72 | Female | Never | Bronchus |
| 4 | Seoul St. Mary's Hospital | 73 | Male | Ex | Bronchus |
| 5 | Seoul St. Mary's Hospital | 67 | Female | Never | Bronchus |
| 6 | Seoul St. Mary's Hospital | 54 | Female | Ex | Bronchus |
| 7 | Seoul St. Mary's Hospital | 59 | Female | Never | Bronchus |
| 8 | Seoul St. Mary's Hospital | 76 | Male | Ex | Bronchus |
| 9 | Seoul St. Mary's Hospital | 73 | Female | Never | Bronchus |
| 10 | Seoul St. Mary's Hospital | 66 | Male | Ex | Bronchus |

**Tissue collection and digestion**

All tracheal or bronchial tissues were processed as previously described [1]. Briefly, tracheal and bronchial tissues were digested with Dispase at 37°C for 1 h. Then, the epithelia were peeled off, digested with trypsin (Sigma-Aldrich) at 37°C for 5 min, and neutralized with blocking buffer (DPBS containing 2% FBS). After centrifugation, pellets were filtered through 40 μm cell strainer, and cells were counted and seeded accordingly. Cow lung tissues were finely minced with razors, incubated in DMEM containing collagenase/hyaluronidase (Stemcell Technologies, #07912) at 37°C for 2 h, and then neutralized with blocking buffer [2]. Then, lung cell pellets were processed in the same way as tracheal tissue preparation as described above.

**Organoid culture**

Media containing 1,000 to 10,000 tracheal epithelial cells from each animal were mixed with growth factor-reduced matrigel (Corning) at 1:1 ratio to a final volume of 100 μl. The mixture was seeded on the apical side of cell culture inserts for 24-well plates. When the seeded mixture solidified, 400 μl of growth medium was added to the basolateral side of the well. The composition of the growth medium varied depending on the species and tissue types. For mouse tracheal organoids, MTEC medium was used [3].

For human, dog, cow, and pig airway organoids, AO medium was used as a base medium [4] and optimized through component addition or removal experiments as displayed in figure 1. All organoids were grown at 37°C in humidified incubator with 5% $CO_2$. The growth media were changed every other day.

**Biosafety**

All experiments involving viral agents, including human influenza and OC43 viruses, were conducted under biosafety level 2 (BSL-2) conditions at DGIST. Manipulations involving infectious virus samples were performed within a Class II biosafety cabinet (BSC) that was airflow-certified and decontaminated with 70% ethanol and ultraviolet (UV) light prior to use. A plastic container containing 70% ethanol was placed inside the BSC before beginning each experiment. Personnel wore appropriate personal protective equipment (PPE), including nitrile gloves, laboratory coats, and eye protection, at all times while handling infectious materials. Upon completion of experiments, all waste materials were soaked in 70% ethanol, collected in biohazard waste bags, and subsequently autoclaved at 125 °C and 20 psi for 1 hr.

**Virus infection**

H1N1 virus (A/human/Korea/KUMC-33/2006/H1N1) was obtained from Korea Bank for Pathogenic Viruses (KBPV). Human coronavirus-OC43 (HCoV-OC43) was obtained from KBPV. SARS-CoV-2 pseudo-entry virus, expressing spike protein, was generated using a SARS-Related Coronavirus 2, Wuhan-Hu-1 Spike-Pseudotyped Lentiviral Kit (BEI Resources). In brief, pseudotyped lentiviral particles containing the spike (S) glycoprotein and green fluorescent protein were produced by transfecting HEK293T cells using Lipofectamine 3000 (Thermo Fisher). The virus-containing medium was harvested at 24 h, 48 h, and 72 h after the transfection, concentrated via ultracentrifugation and titrated by qPCR. Organoids at the size of about 100 μm within passage 0 to 3 were transduced with various viruses in serum-free medium for 2 h at MOI = 1, as indicated in the manuscript and corresponding figure legends. Then, organoids were washed 3 times with 2% FBS/DPBS and cultured for 2 days. Viral production level was measured by a quantitative analysis of viral gene copy number by qPCR with the collected culture medium at 2, 24, and 48 h after viral infection. Organoids were also collected for immunostaining and/or qPCR.

**qPCR**

RNA was extracted using a Quick-RNA Microprep Kit (Zymo Research), and cDNA was synthesized by using a High-Capacity cDNA Reverse Transcription Kit (Thermo) following manufacturer's protocols. qPCR was performed with a StepOnePlus Real-Time PCR system using SYBR Green PCR Master Mix (Thermo). Gene expression was quantified using ΔΔCt method. An absolute viral copy number was quantified using a standard curve for each virus. All primers used are listed in the supplementary table S5.

**Supplementary table S5. Primer sequences for quantitative PCR.**

| Target | | Forward | Reverse |
|---|---|---|---|
| H1N1 | | ATGGTTCTGGCCAGCACTAC | CCATTTGCCTGGCCTGACTA |
| Human coronavirus-OC43 [5] | | AGCAACCAGGCTGATGTCAATACC | AGCAGACCTTCCTGAGCCTTCAA |
| CXCL10 | Human | CTCCAGTCTCAGCACCATGA | GCAGGTACAGCGTACAGTTC |
| | Mouse | GCCGTCATTTTCTGCCTCAT | GATAGGCTCGCAGGGATGAT |
| | Dog | ACCTCTCTCTAGAACTATACGCT | ACACGATGGACTTGCAGGAA |
| | Cow | ACCAAGCCAGTCACCATATCA | CTGGGACTTAGCACATTGACTG |
| HERC5 | Human | CTGGGCCACACTGAGAGTAA | AGTAAACAGCAGCCCATCCT |
| | Mouse | TCGCTCCTACTGTTCAGCAA | CTCACCAGGTCAACATGCAG |
| | Dog | GAGTGACCCAGGTAGCATGT | CCGTTTCCCAGTTGTCCTTC |
| | Cow | GAAGGACAACTGGGAAACGG | TGATTCCCTCCAGCAACCAT |
| OASL | Human | GCGTTTCTGAGCTGTTTCCA | GGACTCTCTGCTCCATCCTC |
| | Mouse | CAGCCTAGTTGCCTCTCCTT | TGTCTGGCTGTACACGACTT |
| | Dog | TCCAACACTCGCATCTGTCT | CTTTGGCGTCAGTGGTGTAG |
| | Cow | TTTCTGTGGTGTCTGAGCCA | CGCACAGGGAGATGTTTCAG |
| IFIT2 | Human | TATCACATGGGCCGACTCTC | ACTTTAACCGTGTCCACCCT |
| | Mouse | ATCACATGGGCCAGTTCTCA | CCTCACAGTCAAGAGCAGGA |
| | Dog | TGCCAAACAATGCCTACCTG | GCTTGTCCCATGGCTTCTTC |
| | Cow | CCTGAGCTGGACTGTGAAGA | CTCTGGGTTCTTGGGGTTCT |
| CCL5 | Human | TATTCCTCGGACACCACACC | ACACACTTGGCGGTTCTTTC |
| | Mouse | TGCCAACCCAGAGAAGAAGT | AGAGCAAGCAATGACAGGGA |
| | Dog | CAGCAGTCGTCTTTGTCACC | CAGGTTCCAGATGCCCTACA |
| | Cow | CACCCACGTCCAGGAGTATT | CCACCCTAGCTCAACTCCAA |
| RSAD2 | Human | ACCCTTCCAAGTCCATCCTG | TCGTTCCACTTTCCGCTCTA |
| | Mouse | ATTCACGCCCAGCTATCCTT | GCTGGTTCTTAGGGGTCACT |
| | Dog | ACCCACATCCTCTGAGTTGG | ATGGGGAGCGGTGATGTAAA |
| | Cow | TCATCGCATGTGGAATCTGC | ACAGGAGCCACAGTAGACAC |
| IFIT1 | Human | GCCTTGCTGAAGTGTGGAGGAA | ATCCAGGCGATAGGCAGAGATC |
| | Mouse | TACAGGCTGGAGTGTGCTGAGA | CTCCACTTTCAGAGCCTTCGCA |

| | | | |
|---|---|---|---|
| | Dog | GAAGGGCCAAAACCAGGAAG | ATAGTTGCCCCATGTGACCA |
| | Cow | AGAAGCAGGTGACCACAGAA | TTCCAGAAATCGGCCGTAGT |
| IRF7 | Human | CGAGCTGCACGTTCCTATAC | AGCAGTTCCTCCGTGTAGC |
| | Mouse | CGGTGATCTTTCCCAGTCCT | TCCCAGTACACCTTGCACTT |
| | Dog | GACCCCACTGACCCACATAA | ATCAGTGGTGGGGTGTCTTC |
| | Cow | TCAAAACCTGTGACGCTTCG | GAAGAGAGAGGAGGGCTGTC |
| IL1B | Human | CCACAGACCTTCCAGGAGAATG | GTGCAGTTCAGTGATCGTACAGG |
| | Mouse | TGGACCTTCCAGGATGAGGACA | GTTCATCTCGGAGCCTGTAGTG |
| | Dog | AAGCCACAAATACCTGGTGC | TTTCATCCCCGTGCACAAAG |
| | Cow | ACCCCAAAGTCTACCCCAAG | ACGGGCCTTTCTTCGATTTG |
| OAS1 | Human | AGGAAAGGTGCTTCCGAGGTAG | GGACTGAGGAAGACAACCAGGT |
| | Mouse | CTGTGCTGACCTCAGAGAAGTC | TGCCCTTGAGTGTGGTGCCTTT |
| | Dog | AGTTGTCAAGGGTGGCTCTT | TCTCGAGCTGCTCCTGAAAA |
| | Cow | GTCCTGAAGCAGCGAAAGAG | CTGGGAGTACTTAGCCAGGG |
| TNFα | Human | TCTGGGCAGGTCTACTTTGG | TGAGCCAGAAGAGGTTGAGG |
| | Mouse | GTGCCTATGTCTCAGCCTCT | CTGATGAGAGGGAGGCCATT |
| | Dog | TCTCGAACCCCAAGTGACAA | CCCATCTGACGGCACTATCA |
| | Cow | TTCAGGAGGTCAAGGTGTCC | AGCAAAAGGAGGCACAAAGG |
| MX1 | Human | CCAGAGGCAGGAGACAATCA | CAGGCTTCGTCAAGATTCCG |
| | Mouse | TGGACATTGCTACCACAGAGGC | TTGCCTTCAGCACCTCTGTCCA |
| | Dog | ACCTCGTCTGTCACCTGAAG | TTCTCCTTCTGCAGAGCCTC |
| | Cow | TGCTAACGTGGACATCGCTA | CCAGATCGGGCTTTGTCAAG |
| MX2 | Human | GCTCCGAGAGAATGGTGACT | TTGGTAGCGGTCTCACTCTG |
| | Mouse | ACCAGAGTTCAGGGAAGAGC | ACTTTGCCTCTCCACTCCTC |
| | Dog | CCCGATCTGACCCTCATTGA | CCACCAGGTTGATCGTCTCT |
| | Cow | AAGTGGAGTGGGAGATTCGG | CCAGGAAGGTCAATGAGGGT |
| IFNγ | Human | TATGCCTGCAATCTGAGCCA | TGGGTACAGTCACAGTTGTCA |
| | Mouse | GATTGCGGGGTTGTATCTGG | GGTCACTGCAGCTCTGAATG |
| | Dog | CACCAGTAAGAGGGAGGACT | TCCTTTTCCGCTTCCTTAGGT |
| | Cow | CTCTGTGGGCTTTTGGGTTT | GCCCACCCTTAGCTACATCT |
| STAT1 | Human | ACCTGCTCCCTCTCTGGAAT | TGAATGTGATGGCCCCTTCC |
| | Mouse | TGGGCGTCTATCCTGTGGTA | TGCGTTCAGACCTCTCTTGG |
| | Dog | CCTGAGGAGTTTGACGAGGT | AGAGTAGCAGGAGGGAGTCA |
| | Cow | TCTTCCTGAACCCACCTTGT | TGTTCAGCTGGTCCACATTG |
| IFNα | Human | GCCATCTCTGTCCTCCATGA | CTGGTAGAGTTCGGTGCAGA |
| | Mouse | CTGCTGGCTGTGAGGAAATAC | CTCTCAGTCTTCCCAGCACA |
| | Dog | GTCCACGTGATGACCCAGAA | AGGTCATCCAGCTGCTCAGA |
| | Cow | TTCACAGAGTCACCCACCTC | GGCAACCCAGAGAGCAGAT |
| TLR3 | Human | GCACAAAGAATGAGGCGACA | CCTGTAACCCGGCTTTTGAG |
| | Mouse | TTGCGTTGCGAAGTGAAGAA | TCAGTTGGGCGTTGTTCAAG |
| | Dog | GATGAGCTGTTGGAGGGTCT | TTGAACACCTCTGCTGGGAT |
| | Cow | ACCTAGAGCTGACCACCAAC | CAAGACCCTTCAGCAACTCG |
| STAT2 | Human | TCTGGTGGAGCAACGTTCAG | TGCTTCAGACCCTGGTAGGT |
| | Mouse | GGACGAAGCTTTTGGGTGTT | TCCTGTTTGAGCTCCAGAGG |
| | Dog | GGGAACGTGCATCCTTCAGA | TGGCAGGAGGCTATCTGAGT |
| | Cow | GCCTTGACTCAGACCAGCTA | ATCTTGTCGAGCCATGTCCA |
| IFNβ | Human | TGCTCTCCTGTTGTGCTTCT | AAGCCTCCCATTCAATTGCC |
| | Mouse | GGTCCGAGCAGAGATCTTCA | CCTGCAACCACCACTCATTC |
| | Dog | TCCACTGGCAGAAGGAACAT | ATGCTGTACTCCTTGGCCTT |
| | Cow | CTGGGGCAGTTACCTTCAAC | AGAATGCCGAAGATGTGCTG |
| IFI44L | Human | ATGGGCACGTTAGGTTGTTG | TGGGCCTGCATACCTCATAG |
| | Mouse | TCAGGGATCCACAATGACAGT | CACGCACAAGTACATGGCTT |
| | Dog | AGGTAAGGCTGTGCTCCAAT | CTTCTGGCTCTCCCCTTCAA |

| | | | |
|---|---|---|---|
| | Cow | TCTAATGATCCCGTGCCTGA | ATGGGGTCACACAGAGTTGG |
| ACTB | Human | CCCTGGACTTCGAGCAAGAG | ACTCCATGCCCAGGAAGGAA |
| | Mouse | GTCGTACCACAGGCATTGTGATGG | GCAATGCCTGGGTACATGGTGG |
| | Dog | CCTCTATGCCAACACAGTGC | CACACAGAGTACTTGCGCTC |
| | Cow | CCCTGGAGAAGAGCTACGAG | TAGTTTCGTGAATGCCGCAG |

**Histology and immunostaining**

*Tissue paraffin embedding and sectioning*

Tracheal or lung tissues were fixed in 4% paraformaldehyde (PFA) at room temperature for 2 h and washed once with 100 mM glycine and subsequently once with PBS. The fixed tissues were dehydrated sequentially in 50%, 70%, 90%, 95%, and 100% ethanol, and then with xylene. Dehydrated tissues were embedded in paraffin, sliced by microtome at a thickness of 8 μm, and directly mounted onto a slide. The sliced tissues were deparaffinized and rehydrated sequentially in xylene, 100%, 95%, 90%, 70%, and 50% ethanol, and water.

*Organoid cryosectioning*

Organoids were fixed by adding an equal volume of warm fixative (2% PFA in 15% sucrose), followed by incubation for 30 min at room temperature (RT). The top 50% of the solution was removed, 2 volumes of fixative were added to the samples, and samples were incubated overnight at 4°C. Then, the samples were washed 3 times with 1x PBS for 10 min each. After the final wash, an equal volume of 30% sucrose was added to the samples, and the samples were incubated for 30 min at RT. The top 50% of the solution was removed, and an equal volume of 30% sucrose was added to the samples. The samples were kept at 4°C until cryosectioning. Before cryosectioning, the samples in 30% sucrose were transferred to a plastic plate and the sucrose solution was carefully removed. The samples were transferred to a plastic mold using a large bore p200 pipette tip and then embedded in OCT (Sakura, #4583). The organoids were positioned to the bottom of the plastic mold using a sharp tungsten needle, frozen in dry ice, and cryosectioned to a thickness of 10 μm using a Cryostat (Leica, CM3050S)

*H&E staining*

Rehydrated tissue samples were stained with hematoxylin and eosin and dehydrated sequentially in 95%, 100% ethanol, and xylene. Stained samples were mounted using a xylene-base mounting medium (Sigma-Aldrich).

*Immunostaining*

Antigens in the tracheal or bronchial samples were retrieved using 1x pH6.1 or pH9 antigen retrieval buffer (Dako, S169984-2 and S236784-2) heated to 90°C for 15 min. Then, samples were cooled down at RT for 2 h. Samples were blocked in 0.1% triton X-100, 3% BSA, 5% normal donkey serum, and 5% normal goat serum in PBS at RT for 1 h. Primary antibody, diluted in antibody diluent (Dako, S080983-2), was added to the samples and incubated overnight at 4°C. Samples were washed once with 0.1% triton X-100 in PBS and incubated with secondary antibody in PBS for 2 h at RT. Then, samples were washed once in PBS, treated with DAPI in PBS for 5 min at RT, and rinsed with tap water. Samples were mounted using a mounting medium (Vector Labs, H-1700) and imaged using an epifluorescence microscope (ZEISS, Axio Vert.A1) or a confocal microscope (ZEISS, LSM800). All antibodies used in this study and their information are listed in the supplementary table S6.

**Supplementary table S6. Antibody information**

| Name | Dilution | Company | Catalogue |
|---|---|---|---|
| Anti-Prosurfactant Protein C (proSP-C) Antibody | 1:300 | Sigma | AB3786 |
| Aquaporin 5/AQP5 Antibody (D-7) | 1:300 | Santa Cruz | sc-514022 |
| Purified anti-Keratin 5 Antibody | 1:100 | Biolegend | 905504 |
| Anti-Acetylated Tubulin antibody, Mouse monoclonal | 1:1000 | Sigma | T7451 |
| CC10 Antibody (T-18) | 1:200 | Santa Cruz | sc-9772 |
| Mouse pIgR Antibody | 1:300 | R&D | AF2800 |
| Anti-Influenza A Virus Nucleoprotein antibody [AA5H] | 1:200 | Abcam | Ab20343 |
| SARS/SARS-CoV-2 Spike Protein S2 Monoclonal Antibody (1A9) | 1:500 | Invitrogen | MA5-35946 |
| Anti-Coronavirus Antibody, OC-43 strain, clone 541-8F | 1:1000 | Sigma | MAB9012 |
| Goat anti-Mouse IgG (H+L) Cross-Absorbed Secondary Antibody, Alexa Fluor™ 488 | 1:1000 | Invitrogen | A11001 |
| Goat anti-Mouse IgG (H+L) Highly Cross-Absorbed Secondary Antibody, Alexa Fluor™ 594 | 1:1000 | Invitrogen | A11032 |
| Goat Anti-Rabbit IgG H&L (Alexa Fluor® 488) | 1:1000 | Abcam | ab150077 |
| Goat Anti-Rabbit IgG H&L (Alexa Fluor® 594) | 1:1000 | Abcam | ab150080 |
| Donkey anti-Goat IgG (H+L) Cross-Absorbed Secondary Antibody, Alexa Fluor™ 488 | 1:1000 | Invitrogen | A-11055 |

| | | | |
|---|---|---|---|
| Donkey anti-Goat IgG (H+L) Cross-Absorbed Secondary Antibody, Alexa Fluor™ 594 | 1:1000 | Invitrogen | A-11058 |

**RNAscope**

RNAscope multiplex fluorescent *in situ* hybridization was performed on FFPE (formalin-fixed, paraffin-embedded) and fixed frozen samples according to the manufacturer's instructions (RNAscope Multiplex Fluorescent Reagent Kit v2; ACDBio) with slight modifications. For FFPE tissue samples, the boiling time in the target retrieval step was reduced from 15 min to 5 min. For fixed frozen samples, the protocol was modified based on samples. For airway organoids of human, mouse, and cow, the samples underwent a reduced target retrieval time from 15 min to 5 min. In the case of dog and pig tracheal organoids, the post-fixation step and the target retrieval step were omitted, and the samples were treated with a reduced amount of proteases: Protease III, diluted 1/15 in PBS (WELGENE, Cat. No. ML 008-02) at 40℃ for 10 min. After probe hybridization, all samples were incubated overnight in 5x SSC (Invitrogen), followed by signal amplification steps performed according to the manufacturer's protocol. Counterstaining was performed with DAPI (Thermo), diluted 1/1,000 in PBS for 2 min at RT. The samples were mounted using Prolong Diamond Antifade Mountant (Invitrogen) and left to cure overnight at RT in a dark environment. The probes and fluorescent dyes used for each species are listed in the supplementary table S7.

**Supplementary table S7. RNAscope probe information.**

| Species | Tissue / Organoid | Probe | Catalog Number | Fluorescent Dye | Catalog Number | Dilution Ratio |
|---|---|---|---|---|---|---|
| Human | Bronchial tissue & organoid | Hs-FOXJ1 | ACDBio, Cat. No. 430921 | Opal 520 | Akoya Biosciences, Cat. No. FP1487001KT | 1/1500 |
| | | Hs-SCGB1A1-C2 | ACDBio, Cat. No. 469971-C2 | Opal 570 | Akoya Biosciences, Cat. No. FP1488001KT | 1/1500 |
| | | Hs-KRT5-O1-C3 | ACDBio, Cat. No. 547901-C3 | Opal 690 | Akoya Biosciences, Cat. No. FP1497001KT | 1/1500 |
| Mouse | Tracheal tissue & organoid | Mm-Foxj1 | ACDBio, Cat. No. 317091 | Opal 520 | Akoya Biosciences, Cat. No. FP1487001KT | 1/1500 |
| | | Mm-Krt5-C2 | ACDBio, Cat. No. 415041-C2 | Opal 570 | Akoya Biosciences, Cat. No. FP1488001KT | 1/1500 |
| | | Mm-Scgb1a1-C3 | ACDBio, Cat. No. 420351-C3 | Opal 690 | Akoya Biosciences, Cat. No. FP1497001KT | 1/1500 |
| Dog | Tracheal tissue & organoid | Cl-FoxJ1 | ACDBio, Cat. No. 1237991-C1 | Opal 520 | Akoya Biosciences, Cat. No. FP1487001KT | 1/2000 |
| | | Cl-Krt5-C2 | ACDBio, Cat. No. 1238001-C2 | TSA Vivid 570 | ACDBio, Cat. No. 323272 | 1/5000 |

| | | | | | | |
|---|---|---|---|---|---|---|
| | | Cl-Scgb1a1-C3 | ACDBio, Cat. No. 1238011-C3 | TSA Vivid 650 | ACDBio, Cat. No. 323273 | 1/5000 |
| Pig | Tracheal tissue | Ss-FoxJ1 | ACDBio, Cat. No. 1238021-C1 | Opal 520 | Akoya Biosciences, Cat. No. FP1487001KT | 1/1500 |
| | | Ss-Krt5-C2 | ACDBio, Cat. No. 1238031-C2 | Opal 570 | Akoya Biosciences, Cat. No. FP1488001KT | 1/1500 |
| | | Ss-Scgb1a1-C3 | ACDBio, Cat. No. 1238041-C3 | Opal 690 | Akoya Biosciences, Cat. No. FP1497001KT | 1/1500 |
| | Tracheal organoid | Ss-FoxJ1 | ACDBio, Cat. No. 1238021-C1 | Opal 520 | Akoya Biosciences, Cat. No. FP1487001KT | 1/2000 |
| | | Ss-Krt5-C2 | ACDBio, Cat. No. 1238031-C2 | TSA Vivid 570 | ACDBio, Cat. No. 323272 | 1/5000 |
| | | Ss-Scgb1a1-C3 | ACDBio, Cat. No. 1238041-C3 | TSA Vivid 650 | ACDBio, Cat. No. 323273 | 1/5000 |
| Cow | Tracheal tissue & organoid | Bt-FoxJ1 | ACDBio, Cat. No. 1237961-C1 | Opal 520 | Akoya Biosciences, Cat. No. FP1487001KT | 1/1500 |
| | | Bt-Krt5-C2 | ACDBio, Cat. No. 1237971-C2 | Opal 570 | Akoya Biosciences, Cat. No. FP1488001KT | 1/1500 |
| | | Bt-Scgb1a1-C3 | ACDBio, Cat. No. 1237981-C3 | Opal 690 | Akoya Biosciences, Cat. No. FP1497001KT | 1/1500 |

**Image acquisition**

The images were taken in a confocal microscope (Zeiss, LSM 800).

**mRNA-sequencing**

RNA was isolated using Monarch Total RNA Miniprep Kit (New England BioLabs). Total RNA concentration was calculated by Quant-IT RiboGreen (Invitrogen). To assess the integrity of the total RNA, samples were run on the TapeStation RNA screentape (Agilent). Only high-quality RNA preparations, with RIN greater than 7.0, were used for RNA library construction. Each library was independently prepared with 10 ng of total RNA from each sample by SMARTer ultra-low input RNA Sample Prep Kit (Clontech Laboratories) following the manufacturer’s protocol. In brief, poly-A containing mRNA molecules were purified using poly-T-attached magnetic beads, and the mRNA was fragmented into small pieces using divalent cations under elevated temperature. The cleaved RNA fragments were copied into first strand cDNA using SMARTScribe reverse transcriptase (Clontech Laboratories) and 3’ SMART CDS primers. Second strand cDNA synthesis was performed using DNA Polymerase I, RNase H and dUTP. These cDNA fragments went through an end repair process, the addition of a single ‘A’ base, and then ligation of the Illumina adapters. The products were then purified and enriched with PCR to create the final cDNA library. The libraries were quantified using KAPA Library Quantification kits for

Illumina Sequencing platforms according to the qPCR Quantification Protocol Guide (KAPA BIOSYSTEMS) and qualified using the TapeStation D1000 ScreenTape (Agilent). Indexed libraries were then submitted to Illumina NovaSeq, and the paired-end (2 × 100 bp) sequencing was performed by Macrogen Incorporated.

#### mRNA-seq data processing

All RNA-seq analysis for each type of organoid was performed in parallel with default parameters, unless otherwise mentioned. Sequencing quality was inspected by FastQC (v.0.11.9) [6]. Trimmomatic (v.0.39) [7] was utilized to filter out the bases with low quality and adapter sequences. After downloading reference genome FASTA and GFF annotation files of each species and H1N1 in NCBI RefSeq (assembly version of GRCh38.p13, Dog10K Boxer Tasha, GRCm39, ARS-UCD1.2 and ViralMultiSegProj274766 for human and dog, respectively), the trimmed reads were aligned using STAR aligner (v.2.7.10a) [8].

### QUANTIFICATION AND STATISTICAL ANALYSIS

#### Image quantification

The number and ratio of viral antigen-expressing cells were averaged from three biological replicates with each specimen counted from 5 different representative fields that include more than 5 organoids.

#### Statistical analysis of fluorescence images

Quantification of fluorescent signals of viral antigens and DAPI was performed using ImageJ [9].

#### Statistical analysis of mRNA-seq data

To increase statistic powers to detect DEGs, genes with little expression were removed using filterByExpr function in edgeR R package (v.3.34.1) [10] considering sample group information of control and H1N1 infection (filterByExpr parameter min.count = 10 & group = 2). Raw counts of duplicated gene symbols were averaged per gene. After those preprocessing steps, DEGs were analyzed by DESeq2 R package (v.1.32.0) [11] and determined by |fold change| ≥ 1.5 & Benjamini-Hochberg FDR < 0.05 in nbinomWaldTest. Volcano plots were visualized using EnhancedVolcano R package (v.1.13.2) [12].

**Enrichment analysis of mRNA-seq data**

To identify the role of DEGs from each species, DEGs were analyzed by gost function in gProfiler2 R package (v.0.2.1) [13]. In addition to the functional annotation of DEGs, the Wald statistic significances of all genes in DESeq2 result objects were used as input of FGSEA R package (v.1.18.0) [14] for the gene set enrichment analysis for lists of gene ontology (GO) terms, which are based on msigdbr R package (v.7.5.1) [15]. The enrichment analyses of both DEGs and total genes were implemented, respectively, to species without ortholog mapping, selecting the GO Biological Process (BP) with the number of genes between 10–500 with the significance cutoff as Benjamini-Hochberg FDR < 0.05.

**Ortholog mapping and cross-species analysis of mRNA-seq data**

For the comparison of gene expression between different species' organoids, gene symbols of each species were converted based on gene symbol in human by gorth function in gprofiler2 R package (v.0.2.1) [13]. If multiple ortholog genes were present in one species, only one was chosen as the following priorities: (1) the gene with the same symbol, (2) the gene included in DEGs, (3) the gene showing the maximal |$\log_2$ fold change. To investigate the effect of viral infection common to tropic species, over-representation analysis was implemented with the upregulated DEGs intersecting between human and dogs by gost function in gProfiler2 R package (v.0.2.1) [13]. When comparing the expression pattern between H1N1-infected organoids of each species, raw counts of total 12,481 mapped genes of all 8 control and H1N1-infected organoid samples were processed into a new DESeq2 object. From the DESeq2 object, the gene lengths of each species were used for normalization. Gene lengths were calculated as the sum of exon lengths in each gene, by reading GFF annotation files of each species and H1N1 genomes using the GenomicFeatures R package (v.1.44.2) [16]. Furthermore, the species-specific expression patterns were eliminated with removeBatchEffect function from limma R package (v.3.48.3) [17]. The raw counts were $\log_2$-normalized according to the total library size by rlog function in DESeq2 (rlog parameter blind = FALSE). The heatmap of rlog counts was visualized by pheatmap R package (v.1.0.12) [18].

**Comparison between H1N1 and SARS-CoV-2 of mRNA-seq data**

As an external dataset, the raw counts of GSE150819 [19] were processed with the same DEG and enrichment analysis with DESeq2, gProfiler2, and GSEA to compare gene expression of H1N1-infected human bronchial organoids with that of SARS-CoV-2-infected ones. When comparing the expression pattern of each sample with normalized counts, a new DESeq2 object was produced containing both H1N1 and SARS-CoV-2 datasets with the common 12,803 genes. To remove batch effects between the datasets, removeBatchEffect function from limma R package (v.3.48.3) [17] was applied for the rlog-normalized count containing both H1N1 and SARS-CoV-2 treated samples (rlog parameter blind = FALSE), which was then visualized by pheatmap R package (v.1.0.12) [16].

### Supplementary figures

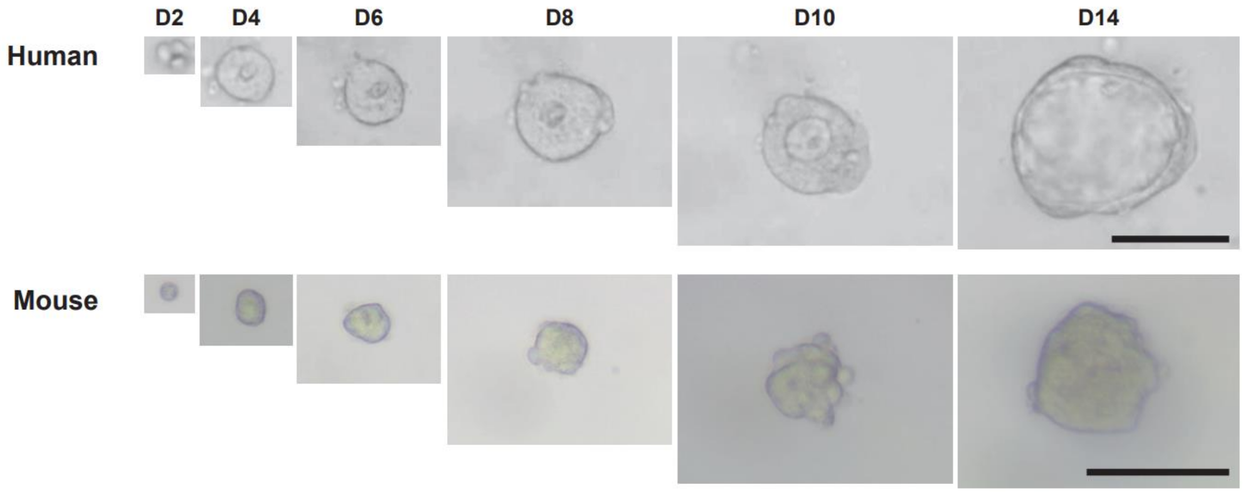


Figure S1. Time lapse images of human and mouse airway organoids. Scale bars: 100 μm.

Figure S2

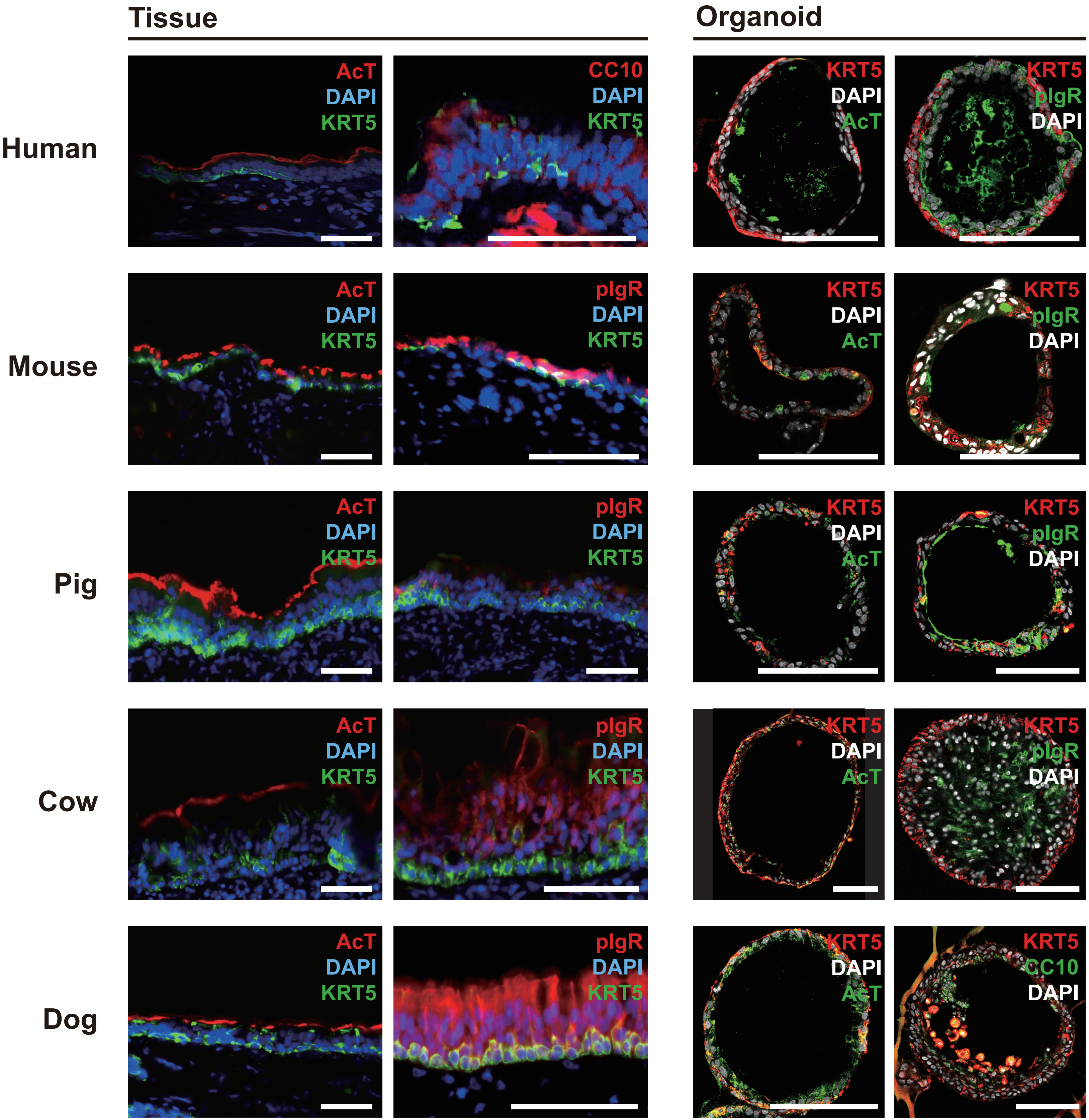

Tissue
Organoid
Human
Mouse
Pig
Cow
Dog
AcT
DAPI
KRT5
CC10
pIgR

Figure S2. Organotypic representation of airway organoids of human, mouse, cow, dog, and pig.

A) Immunofluorescence staining of tracheal or bronchial epithelium. KRT5, pIgR, CC10, and AcT were used as the markers for basal cells, club cells, and ciliated cells, respectively.

B) Immunofluorescence staining of tracheal or bronchial organoids.

Scale bars: 100 μm.

Figure S3

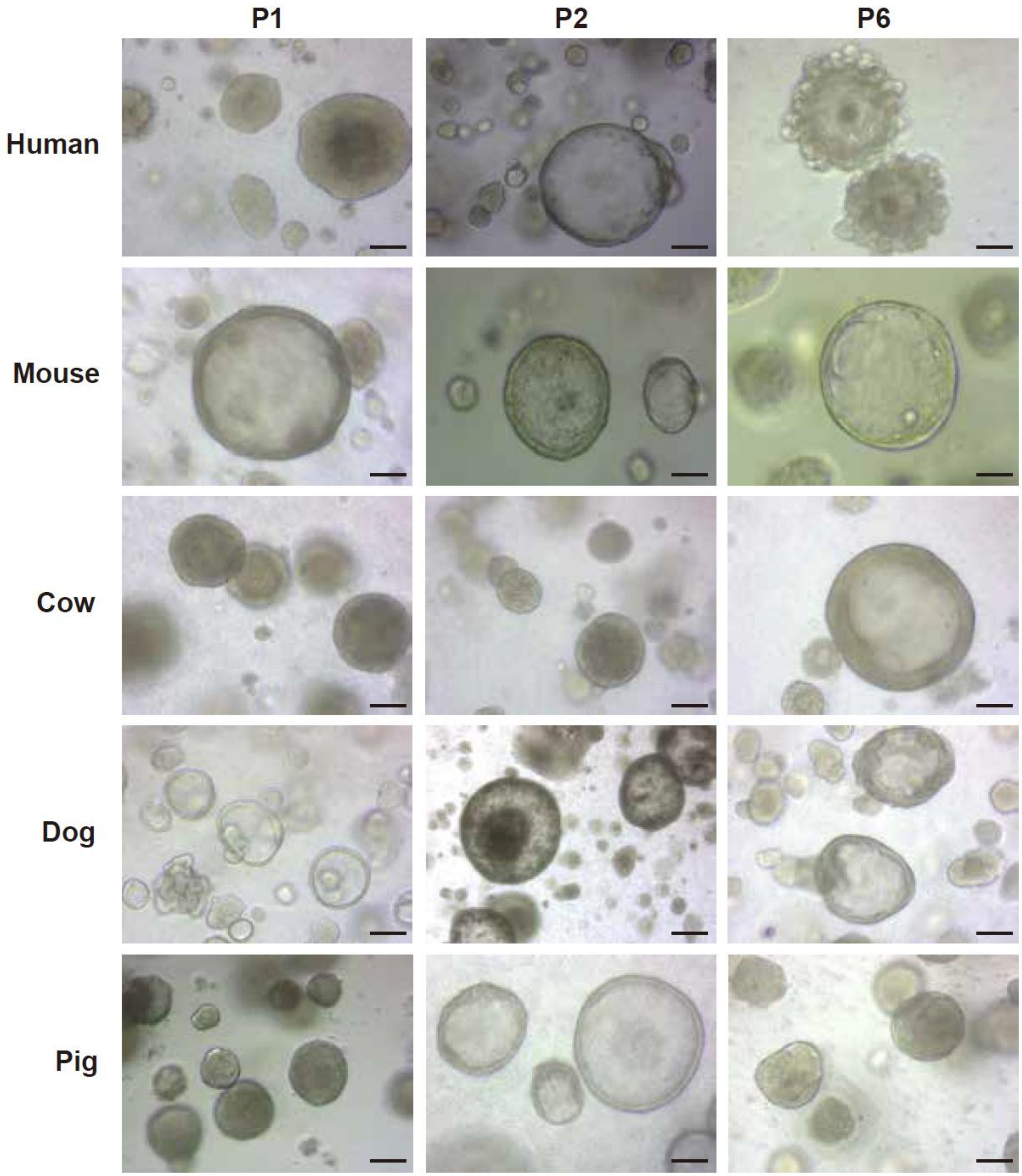


Figure S3. Long-term, 3D cultures of airway organoids of human, mouse, cow, dog, and pig. Bright-field images of airway organoids of passage 1 to 6 are shown. Scale bars: 100 μm.

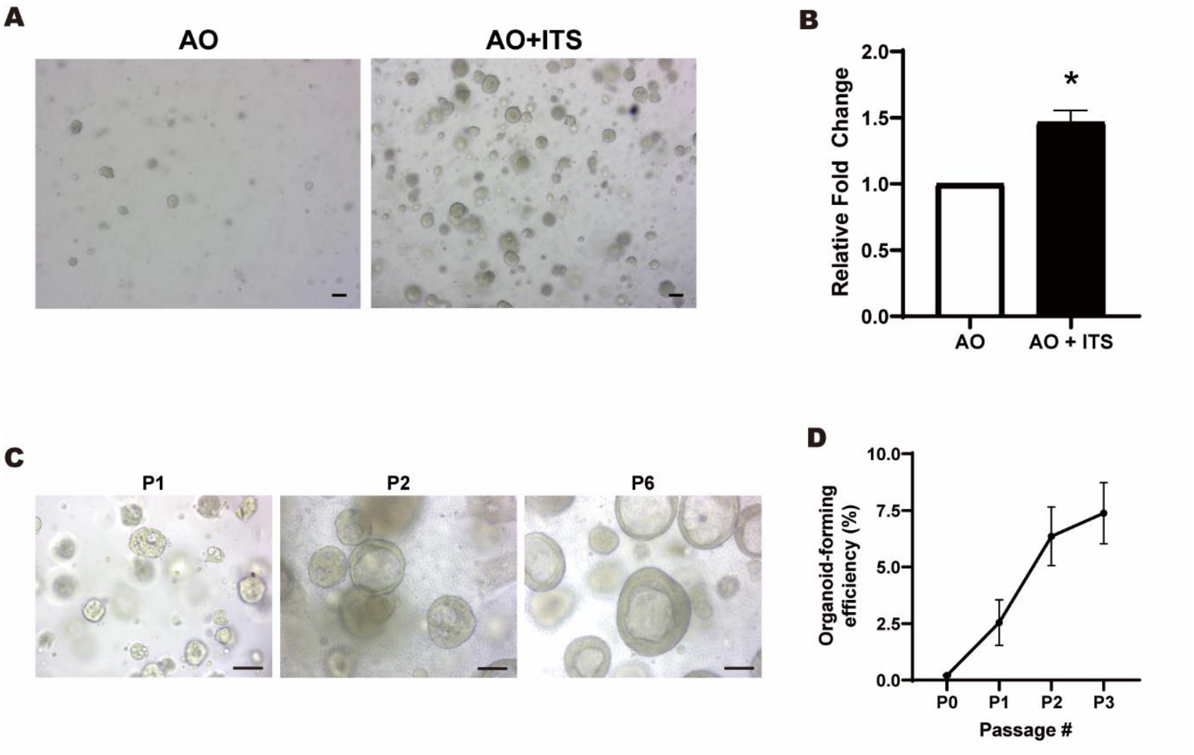


Figure S4. Establishment of cow bronchioalveolar organoids.

A-B) The effect of ITS addition for the culture of cow bronchioalveolar organoids. (A) Representative bright-field organoid images of each culture condition. (B) Relative numbers of organoids in each culture condition.

C) Long-term, 3D cultures of cow bronchioalveolar organoids. Bright-field images of bronchioalveolar organoids of passage 1 to 6 are shown. Scale bars: 100 μm.

D) Organoid-forming efficiency of cow bronchioalveolar organoids along passages (N=3).

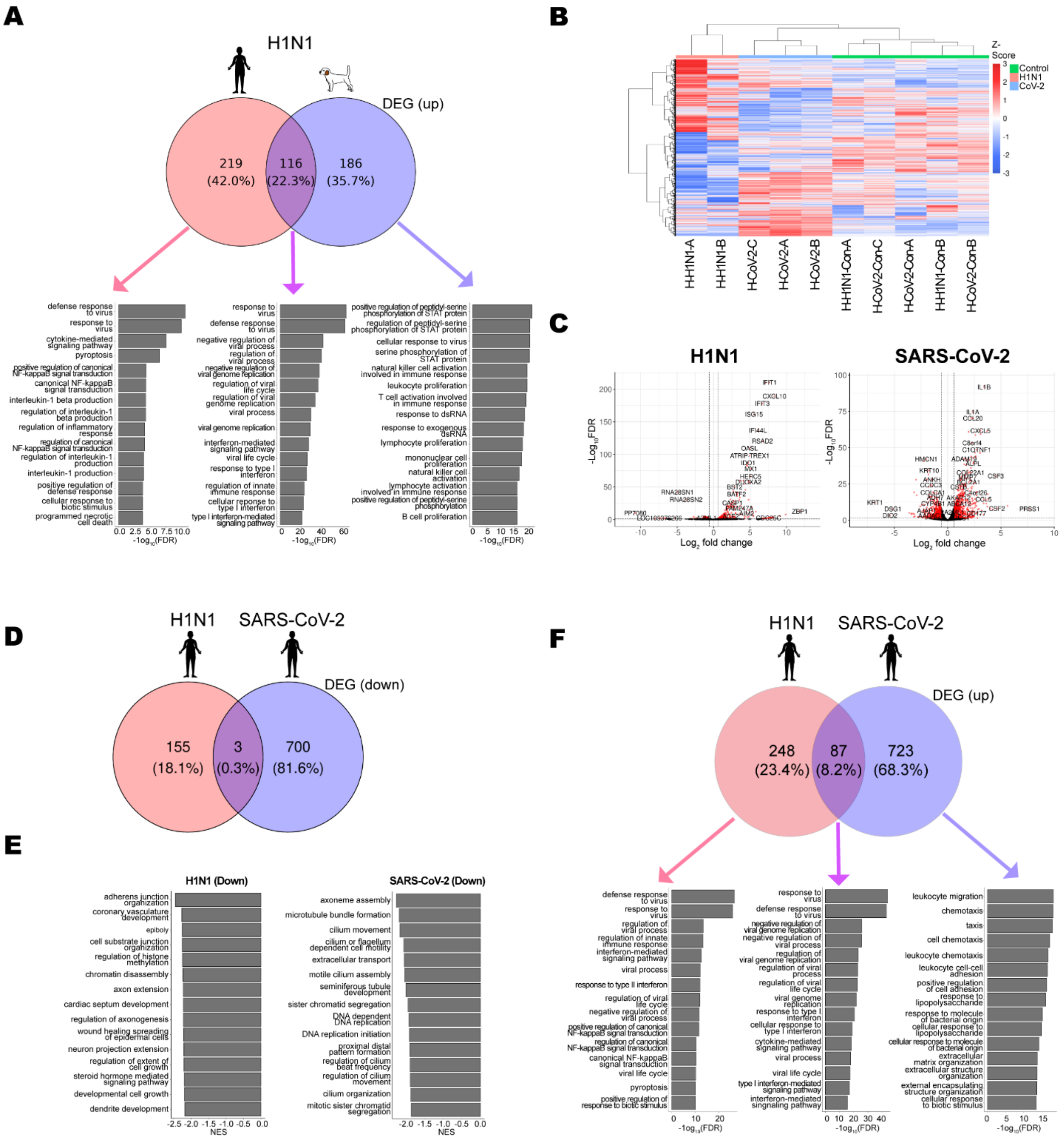

A
H1N1
DEG (up)
219 (42.0%)
116 (22.3%)
186 (35.7%)
B
C
H1N1
SARS-CoV-2
D
H1N1
SARS-CoV-2
DEG (down)
155 (18.1%)
3 (0.3%)
700 (81.6%)
E
H1N1 (Down)
SARS-CoV-2 (Down)
F
H1N1
SARS-CoV-2
DEG (up)
248 (23.4%)
87 (8.2%)
723 (68.3%)

Figure S5. H1N1 and SARS-CoV-2 induce minimal overlap of transcriptomic changes.

A) Bar graph representing GSEA for GO:BP of commonly and species-specifically upregulated genes in H1N1-infected human and dog airway organoids compared to their own mock-infected organoids.

B) Hierarchical clustering of whole transcriptomes from control, H1N1- and SARS-CoV-2 infected human airway organoids. H: human, CoV-2: SARS-CoV-2.

C) Volcano plots showing DEGs between H1N1-infected and mock-infected human airway organoids (left) and SARS-CoV-2-infected and mock-infected ones (right). DEGs (FDR $< 0.05$; $\log_2 FC > \log_2(1.5)$) are indicated in red. Representative genes are denoted by their gene name.

D) Venn diagrams showing down-regulated DEGs in H1N1- and SARS-CoV-2-infected human airway organoids compared to mock-infected organoids.

E) Bar graph representing GSEA downregulated in H1N1- or SARS-CoV-2-infected organoids compared to mock-infected human airway organoids.

F) Bar graph representing GSEA of commonly or virus-specifically upregulated genes in H1N1- or SARS-CoV-2-infected organoids compared to mock-infected human airway organoids.

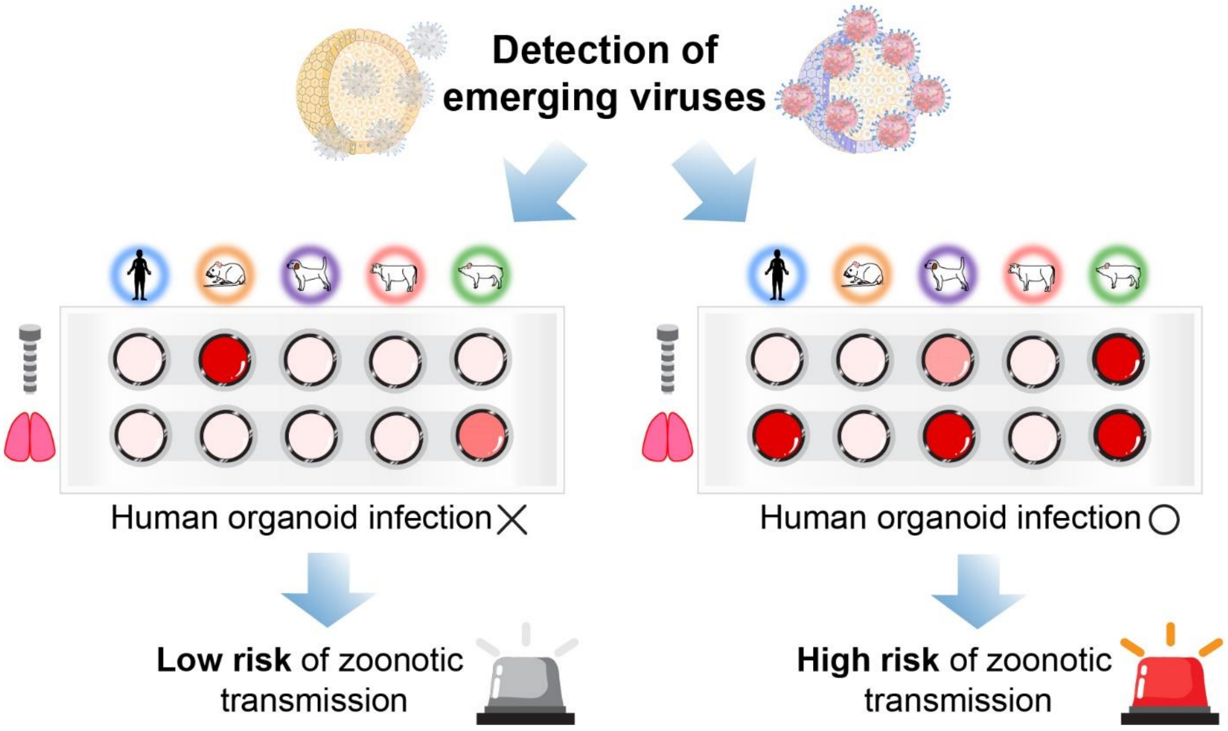


Figure S6. A utility model of a comparative airway organoid platform
